# Lkb1 is a master regulator of VSMC fate and function in mice

**DOI:** 10.1101/2020.12.12.422410

**Authors:** Zhaohua Cai, Ping Song, Shaojin You, Zhixue Liu, Fujie Zhao, Jing Mu, Xiaoxu Zheng, Ye Ding, Lei Xiao, Tharmarajan Ramprasath, Yu Qiu, Ben He, Ming-Hui Zou

## Abstract

Acquisition and maintenance of vascular smooth muscle cell (VSMC) fate are important for vascular development and homeostasis; however, little is known about the key determinant for VSMC fate and vascular homeostasis. We found that VSMC-specific *Lkb1* ablation in *Lkb1*^*flox/flox*^;*Tagln-Cre* mice caused severe vascular abnormalities and embryonic lethality. VSMC-specific deletion of *Lkb1* in tamoxifen-inducible *Lkb1*^*flox/flox*^;*Myh11-Cre/ERT2* mice progressively induced aortic/arterial dilation, aneurysm, rupture, and premature death. Single-cell RNA sequencing and imaging-based lineage tracing showed that *Lkb1*-deficient VSMCs underwent dynamic transcriptional reprogramming and transformed gradually from early modulated VSMCs to fibroblast-like, chondrocyte-like, and even osteocyte-like cells. VSMC transformation followed by extracellular matrix remodeling and inflammatory cell infiltration contributed to the arterial aneurysm formation in tamoxifen-induced *Lkb1*^*flox/flox*^;*Myh11-Cre/ERT2* mice. Finally, we found that VSMC-specific *Lkb1* ablation resulted in decreased vascular contractility, hypotension, and impaired responses to angiotensin II and vessel injury *in vivo. Lkb1* is therefore a key determinant of mouse VSMC fate that prevents VSMC reprogramming and sustains vascular homeostasis. Our findings have important implications for understanding the pathogenesis of aortic aneurysm.

---

The vascular smooth muscle cell (VSMC) is a differentiated cell type located in the medial layer of vessels. Despite being highly specialized, VSMCs retain remarkable phenotypic plasticity. This plasticity has been frequently associated with VSMC phenotypic switching from a contractile to a synthetic phenotype (*1–3*). However, VSMCs can also transdifferentiate/transform into other cell types, including macrophage-like, fibroblast-like, osteochondrocyte-like, or mesenchymal stem-cell–like cells (*4–8*), indicating that this plasticity can extend lineage boundaries. Although emerging evidence strongly implicates the loss of VSMC contractile properties in life-threatening vascular disorders, such as aneurysms, aortic dissection, and even aortic rupture (*9–12*), it remains unclear whether there is a cause-and-effect relationship between VSMC transformation or changes in VSMC fate and arterial dilation or aneurysm development. Moreover, the master regulator for VSMC fate and plasticity remains largely unknown.

Liver kinase b1 (Lkb1), a tumor suppressor, is mutated and strongly implicated in Peutz– Jeghers syndrome, cervical carcinoma, and non-small-cell lung carcinomas (*13–15*). Lkb1 regulates the metabolism of muscle (*16*), liver (*17*), pancreas (*18*), hematopoietic stem cells (*19*), and T_reg_ cells (*20*). Deletion of *Lkb1* in mammalian neurons (*21*), epithelial cells (*22*), and T_reg_ cells (*23*) disrupts their polarity, differentiation, or lineage identity. Mice deficient in *Lkb1* die at mid-gestation with vascular abnormalities, mesenchymal cell death, and neural tube defects (*24*). Deletion of *Lkb1* in endothelial cells causes endothelial dysfunction and hypertension in mice (*25*). However, Lkb1 is not known to regulate VSMC fate and vascular homeostasis.

## Results

### Cardiovascular defects in smooth-muscle-cell–specific *Lkb1* deficient mice

To explore the role of Lkb1 expressed by VSMCs in vessel development and function, we generated mice with smooth muscle cell (SMC)-specific *Lkb1* deletion (*Lkb1*^*SMKO*^) by intercrossing *Lkb1*^*flox/flox*^ mice with *Tagln-Cre* mice. Unexpectedly, the breeding resulted in no viable offspring with the *Lkb1*^*flox/flox*^;*Tagln-Cre*^*tg*^ genotype. Genotyping of 11 stillborn offspring identified 8 with the *Lkb1*^*flox/flox*^;*Tagln-Cre*^*tg*^ genotype and 3 with the *Lkb1*^*flox/+*^;*Tagln-Cre*^*tg*^ genotype (data not shown). Of 80 embryos dissected between embryonic day (E)8.5 and 20.5, we obtained 26 *Lkb1*^*flox/flox*^;*Tagln-Cre*^*tg*^ embryos which were indeed present at the Mendelian ratio, indicating that Lkb1 is dispensable for implantation (table S1). However, of the 26 *Lkb1*^*flox/flox*^;*Tagln-Cre*^*tg*^ embryos, 21 exhibited visible abnormalities, including small size, developmental delay, absence of normal vessel structures, and other defects (table S1 and fig. S1, A to D). Histological and immunofluorescent staining revealed complete absence of vessel structure and VSMC staining in *Lkb1*^*SMKO*^ embryos (fig. S1, E and F). These observations suggest that SMC-specific *Lkb1* deletion resulted in severe vascular abnormalities and embryonic lethality.

Because of the embryonic lethality observed after SMC-specific *Lkb1* ablation, we generated mice with tamoxifen (TAM)-inducible SMC-specific *Lkb1* deletion (*Lkb1*^*SMiKO*^) by intercrossing *Lkb1*^*flox/flox*^ mice with *Myh11-Cre/ERT2* mice. In *Lkb1*^*SMiKO*^ mice, *Lkb1* was ablated in mature VSMCs after TAM induction (Fig. 1A). Remarkably, all *Lkb1*^*SMiKO*^ mice prematurely died within 8 months post-TAM induction, whereas the age-matched wild-type (WT) control mice survived for the duration of the experiment (Fig. 1B). At necropsy, 40 of 42 (95.2%) *Lkb1*^*SMiKO*^ and 0 of 42 WT control mice developed aortic and/or arterial rupture at an average of 173.0 ± 5.5 days post-TAM induction (Fig. 1, C and D). The aortic and/or arterial rupture generally occurred in the abdominal aorta (62.5%), femoral and/or popliteal artery (40.0%), and renal artery (30.0%) in *Lkb1*^*SMiKO*^ mice (fig. S2, A to C). Histopathological analysis revealed altered vessel structure and evidence of massive hemorrhage, confirming aortic/arterial rupture (Fig. 1E). When sacrificed beyond 4.5 months post-TAM induction, all *Lkb1*^*SMiKO*^ mice exhibited aneurysm formation in abdominal aorta, renal artery, iliac artery, femoral artery, and/or popliteal artery (Fig. 1F and fig. S2, B and C). Over the course of 4.5-month follow-up, both the abdominal and ascending thoracic aortas in *Lkb1*^*SMiKO*^ mice showed significant and progressive dilation of their internal diameters compared with those in WT control mice (Fig. 1, G to J). Altogether, these data demonstrate that deletion of *Lkb1* specifically in mature VSMCs gradually caused aortic/arterial dilation, aneurysm, rupture, and premature death, indicating that Lkb1 plays critical roles in maintaining vascular integrity and homeostasis.

**Fig.1.**
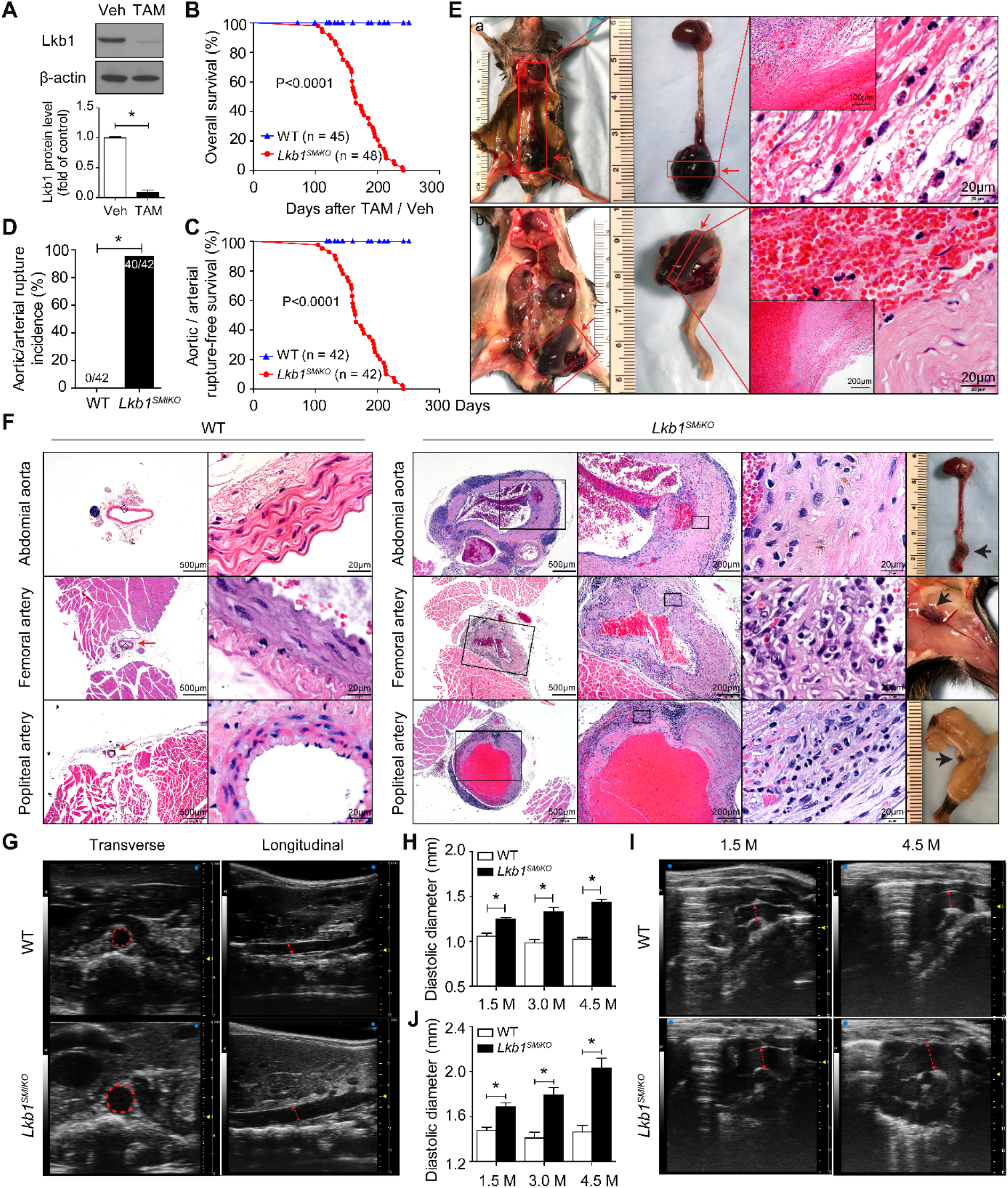
Deletion of *Lkb1* in mature VSMCs causes aortic or arterial rupture and premature death. **(A)** Analysis of Lkb1 expression in primary VSMCs isolated from the aortas of *Lkb1*^*flox/flox*^;*Myh11-Cre/ERT2* mice treated with tamoxifen (TAM) or vehicle (Veh). **(B and C)** Overall survival and aortic/arterial rupture-free survival of wild-type (WT) and *Lkb1*^*SMiKO*^ mice. Statistical analysis was performed using the log-rank test. **(D)** Aortic/arterial rupture incidence in WT and *Lkb1*^*SMiKO*^ mice. Statistics were determined using Fisher’s exact test. **(E)** Macroscopic images and hematoxylin-and-eosin (H&E) staining illustrating ruptured vessels in *Lkb1*^*SMiKO*^ mice. Red arrows denote ruptured aorta (a) or femoral artery (b). **(F)** Macroscopic images and H&E staining of abdominal aorta, femoral artery, and popliteal artery in WT and *Lkb1*^*SMiKO*^ mice. Boxes indicate fields shown at higher magnification in images to the immediate right. Red arrows denote the arteries. Black arrows denote aneurysms. **(G-J)** Representative images and quantification of the abdominal (G and H) and ascending thoracic (I and J) aortas monitored by ultrasound from WT and *Lkb1*^*SMiKO*^ mice. Red dashed circles and arrows indicate aortic internal diameter.

### Single-cell RNA sequencing identifies transformed VSMC populations and increased inflammatory cells in *Lkb1*^*SMiKO*^ aorta

To determine the effect of Lkb1 on VSMCs, we performed a time series of droplet-based single-cell RNA sequencing (scRNA-seq) analyses, based on the single aortic cells isolated from WT and *Lkb1*^*SMiKO*^ mice at 1.0, 3.5, and 4.5 months post-TAM induction (fig. S3A). Data from WT and *Lkb1*^*SMiKO*^ mice were separated into two groups: WT-1M and KO-1M (defined as the early-stage group) and WT-4.5M and KO-3.5M and KO-4.5M (defined as the advanced-stage group). Droplets (n=9202) from early-stage aortas were analyzed, yielding 6928 cells (3154 from WT-1M and 3774 from KO-1M) after rigorous quality control (see the Materials and Methods). Graph-based clustering algorithms were used to define 12 distinct clusters, which were further annotated into 5 major cell types by the canonical marker expression: VSMC (clusters 1, 3, 5, 8, expressing VSMC differentiation markers *Myh11, Cnn1*, and *Acta2)*, early modulated VSMC (clusters 0, 2, 6, 7, 10), fibroblast (cluster 4, expressing *Clec3b* and *Serpinf1)*, endothelial cell (cluster 9, expressing *Vwf* and *Sox18)*, and macrophage (cluster 11, expressing *Ccl4* and *Lyz2)* (Fig. 2A and fig. S3, B and C). Clusters 0, 2, 6, 7, and 10 were annotated as early modulated VSMC based on the following characteristics. First, the cells exhibited expression levels of the VSMC differentiation markers consistent with normal VSMCs. Second, the cells displayed high expression levels of *Tcap* (telethonin) and *Rasl10b* (RAS-family 10 member B), both of which are highly expressed in cardiac and skeletal muscle; *Hapln1* (hyaluronan and proteoglycan link protein 1), an extracellular matrix (ECM) component found in cartilage; and many other genes, including *Ifi27l2a, Rbp4, Por*, and *Egln3*. Third, the cells showed slightly upregulated expression of pro-inflammatory genes, including *H2-Ab1, Cd74, Cxcl12, and Alcam* (Fig. 2A and fig. S3, B and C). We observed that VSMCs from *Lkb1*^*SMiKO*^ mice were exclusively transformed into early modulated VSMCs during this early stage (Fig. 2A).

**Fig.2.**
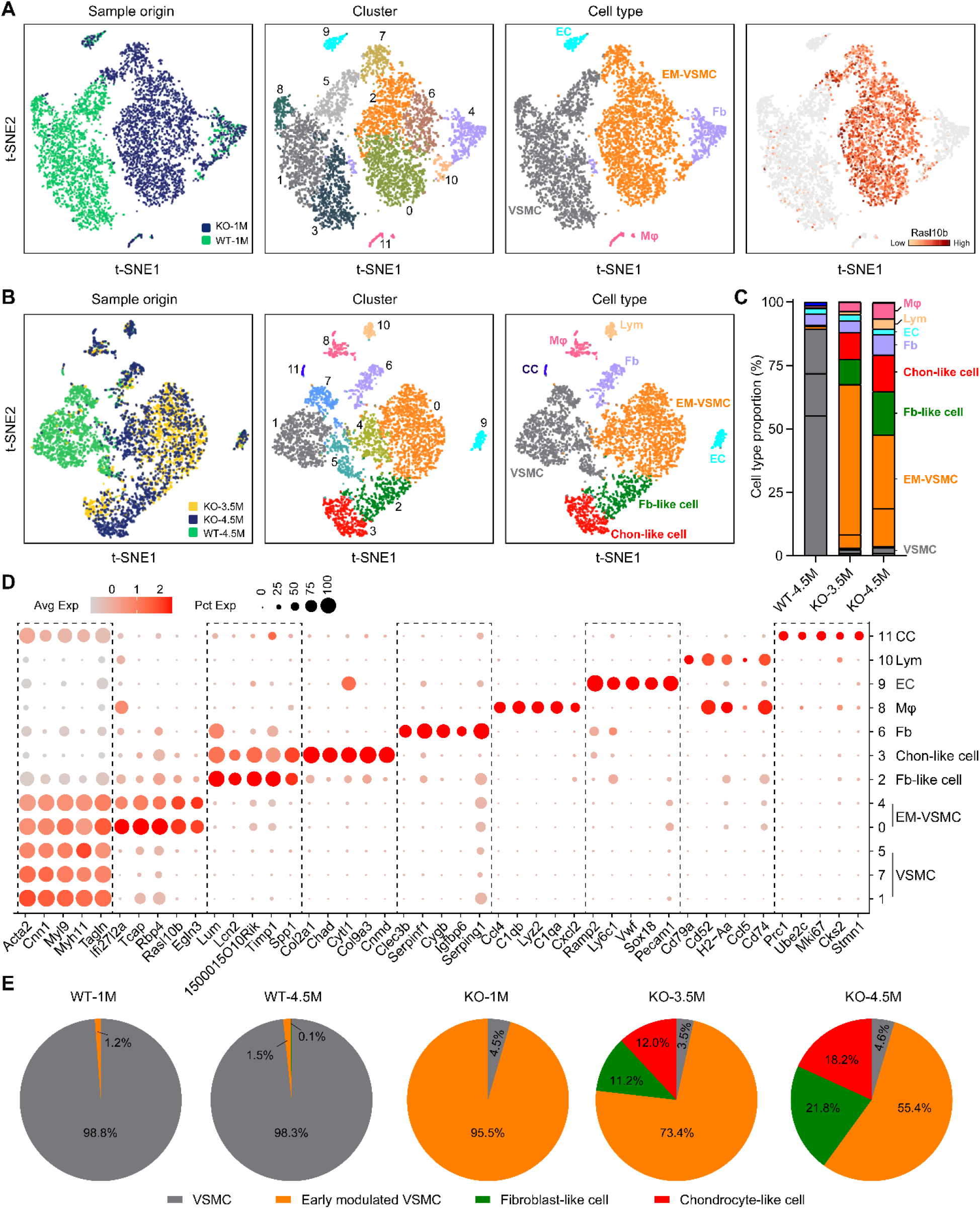
Single-cell RNA-seq identifies transformed cell populations in *Lkb1*^*SMiKO*^ mice. **(A)** Single-cell RNA-seq of the aortas from two independent samples of WT and *Lkb1*^*SMiKO*^ mice at 1.0 month (WT-1M and knockout [KO]-1M, respectively) post-tamoxifen (TAM) induction using the 10x Genomics Chromium platform. *t*-SNE clustering of 6928 single cells (3154 from WT-1M and 3774 from KO-1M) colored by sample origin, cell cluster, and cell type (VSMC, vascular smooth muscle cells; EM-VSMC, early modulated VSMCs; EC, endothelial cell; Fb, Fibroblast; Mφ, macrophage). **(B)** Single-cell RNA-seq of the aortas from three independent samples of WT and *Lkb1*^*SMiKO*^ mice at 3.5 and 4.5 months post-TAM induction. *t*-SNE plots of 5560 single cells (1699 from WT-4.5M, 1642 from KO-3.5M, and 2219 from KO-4.5M), colored by sample origin, cell cluster, and cell type. Lym, lymphocyte; CC, cycling cell; Fb-like cell, fibroblast-like cell; Chon-like cell, chondrocyte-like cell. **(C)** Bar plot showing the relative distribution of each cluster identified in (B) in different samples of WT-4.5M, KO-3.5M, and KO-4.5M. **(D)** Bubble plot for gene expression of five cell-type differentiating markers. Dot color scale indicates the average expression level (Avg Exp) and dot size represents percentage of cells in each cluster that express at least one transcript of each gene (Pct Exp). **(E)** Pie chart showing percentage of each cell type among VSMCs of WT and *Lkb1*^*SMiKO*^ mice at indicated time point post-TAM induction.

Next, we investigated single-cell transcriptomes from advanced-stage aortas. We analyzed 7463 droplets in this stage, yielding 5560 cells (1699 from WT-4.5M, 1642 from KO-3.5M, and 2219 from KO-4.5M) after rigorous quality control (see the Materials and Methods), and identified 12 clusters, which were manually curated into 9 different cell types based on distinct marker expression. The most notable change was that the cells in *Lkb1*^*SMiKO*^ mice (cells in KO-3.5M and KO-4.5M) shifted from clusters 1, 5, 7 to clusters 0, 4, 2, 3. In other words, cells from clusters 1, 5, and 7 were strikingly decreased, whereas cells from clusters 0, 4, 2, and 3 were dramatically increased in *Lkb1*^*SMiKO*^ mice, compared with cells in WT control mice (Fig. 2, B and C). When zooming in on the detailed gene expression profiles of these clusters (Fig. 2D and fig. S4), clusters 1, 5, and 7, which were mainly composed of WT cells showing high-level expression of the VSMC differentiation markers (*Myh11, Acta2*, and *Tagln)* but no expression of pro-inflammatory genes (*H2-Ab1, H2-Eb1*, and *Cd74)*, fibroblast markers (*Lum, Fmod*, and *Lcn2)*, or chondrocyte markers (*Col2a1, Col9a1*, and *Chad)*, suggesting normal VSMCs. Clusters 0 and 4, which were mainly composed of cells from *Lkb1*^*SMiKO*^ mice exhibited expression levels of the VSMC differentiation markers consistent with normal VSMCs and moderately upregulated expression of pro-inflammatory genes, suggesting early transformed VSMCs. Cluster 2, which was mainly composed of cells from *Lkb1*^*SMiKO*^ mice, exhibited dramatically downregulated expression of the VSMC differentiation markers and highly upregulated expression of pro-inflammatory genes (*H2-Ab1, H2-Eb1, Cd74, C3, Cxcl12*, and *Vcam1)* and fibroblast markers, suggesting fibroblast-like cells. Cells in cluster 3 showed exclusively high-level expression of chondrocyte markers and no expression of the VSMC differentiation markers, suggesting chondrocyte-like cells.

Collectively, we found that normal VSMCs dominated the WT aortic cell population (approximately 90% of aortic cells), but from early to advanced stages, *Lkb1*^*SMiKO*^ VSMCs shifted from early modulated VSMCs to fibroblast-like and chondrocyte-like cells (Fig. 2E). At 4.5 months post-TAM induction, 21.8% and 18.2% of *Lkb1*^*SMiKO*^ VSMCs had transformed into fibroblast-like and chondrocyte-like cells, respectively, accompanied by 55.4% of early modulated VSMCs (Fig. 2E). In addition, infiltration of inflammatory cells, particularly macrophages and lymphocytes, was significantly increased in *Lkb1*^*SMCiKO*^ aorta. These inflammatory cells constitute up to 10% of total aortic cells in *Lkb1*^*SMiKO*^ aorta at 4.5 months post-TAM induction (Fig. 2, B and C).

To more definitively differentiate distinct transformed VSMC populations, we combined the datasets from the early and advanced stages using robust principal component analysis (RPCA) (see the Materials and Methods). Both normal VSMCs and early modulated VSMCs were clustered together in early and advanced stages; however, fibroblast-like and chondrocyte-like cells were exclusively derived from advanced-stage aorta of *Lkb1*^*SMiKO*^ mice (fig. S5, A to C). Taken together, these data suggest that *Lkb1*-deficient mouse VSMCs might undergo a continuous and progressive cell transformation from early modulated VSMCs to fibroblast-like and chondrocyte-like cells. Different *Lkb1*-deficient VSMCs might stay in different stages of cell transformation.

### Single-cell trajectories reveal a gradual cell transformation in *Lkb1*^*SMiKO*^ mice

To further test this hypothesis, we applied the Monocle 2 algorithm to perform cell trajectory analysis using pseudotime reconstitution of four cell groups from the advanced-stage aorta (Fig. 3A). The pseudotime trajectory presented an organized progression of cells from VSMCs to early modulated VSMCs to fibroblast-like and chondrocyte-like cells (Fig. 3, A and B and fig. S6, A and B). Pseudotemporal expression dynamics of specific representative genes also marked the progression of VSMCs to early modulated VSMCs and then fibroblast-like and chondrocyte-like cells (Fig. 3C). VSMC differentiation markers *Myh11* and *Acta2* showed a gradient of decreasing expression along the pseudotime trajectory (Fig. 3D). By contrast, there was a striking and progressive increase in small leucine-rich proteoglycans such as lumican (*Lum)*, fibromodulin (*Fmod)*, and decorin (*Dcn)* (fig. S6C) and chondrocyte differentiation markers, such as *Col2a1, Col9a1, Col27a1, Sox9*, and osteoprotegerin (*Tnfrsf11b)* along the pseudotime trajectory from VSMC to chondrocyte-like cell (Fig. 3D and fig. S6C). Similarly, fibroblast markers were upregulated, including dermatopontin (*Dpt)*, fibulin 1 (*Fbln1)*, and *Tgfbi*, predominantly in fibroblast-like cell clusters (Fig. 3D and fig. S6C). Notably, at the transcription level, VSMCs undergoing cell transformation also appeared to be shifting towards a pro-inflammatory phenotype (Fig. 3D and fig. S6C). The results presented here delineate, for the first time, the progressive transformation of mouse VSMCs in an *in vivo* system.

**Fig.3.**
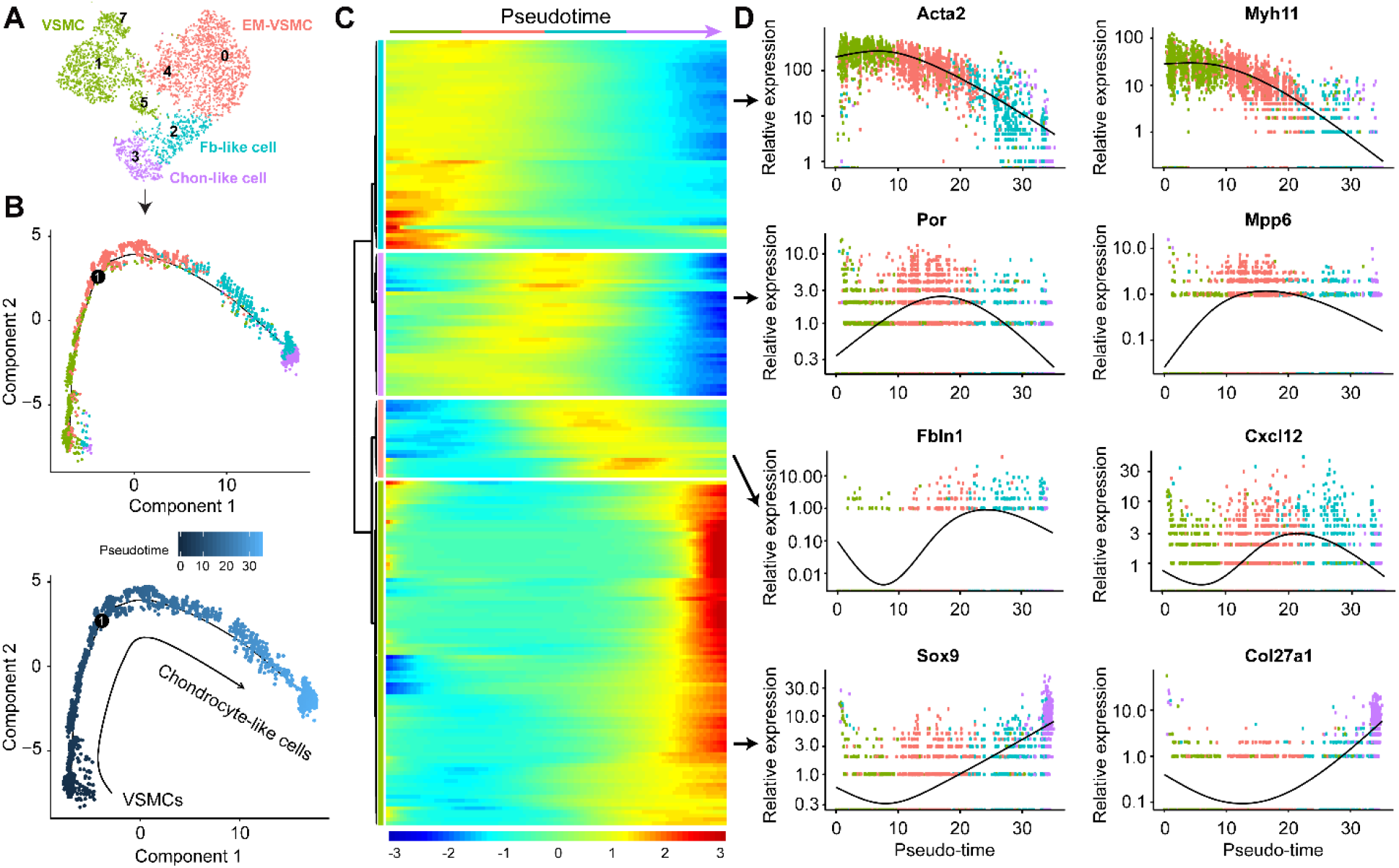
Single-cell trajectory reveals a VSMC transformation pathway, from early modulated VSMCs to fibroblast-like to chondrocyte-like cells in *Lkb1*^*SMiKO*^ mice. **(A-B)** Unsupervised analysis of single-cell gene expression profiles with Monocle revealed a linear trajectory. Cells within specific lineage clusters were selected, visualized using t-SNE visualization (Seurat R package), and then ordered based on a reversed graph embedding method (Monocle 2). **(C)** Heatmap of differentially expressed genes, ordered based on their common kinetics. The Expression Z score indicates changes in a gene relative to its dynamic range over pseudotime. **(D)** Kinetic diagrams showing the expression of some genes over pseudotime.

To obtain molecular insight into the progressive transformation of *Lkb1*^*SMiKO*^ VSMCs, we further performed differential expression analysis focusing on transcription factors. We identified the top 50 transcription factors showing dynamic expression over the pseudotime (fig. S7). Among them, Klf4 progressively increased along the pseudotime trajectory from VSMC to chondrocyte-like cell (fig. S7, A and B). This result is consistent with the well-known role of Klf4 in induction of VSMC transformation (*5, 26, 27*). Additionally, the orphan nuclear receptors Nr4a1, Nr4a2, and Nr4a3 strikingly increased over the pseudotime (fig. S7, A and B), suggesting their important roles in regulation of VSMC fate. However, some transcription factors, such as Tgfb1i1, decreased over the pseudotime, (fig. S7, A and B). Our findings reveal that VSMC transformation in *Lkb1*^*SMiKO*^ mice was driven by a highly dynamic transcriptional reprogramming.

### Visualization of transformed cell populations in *Lkb1*^*SMiKO*^ mice

We then sought to localize and visualize the transformed VSMCs in aortas or arteries of *Lkb1*^*SMiKO*^ mice. Therefore, we first checked the thoracic aortic morphology of *Lkb1*^*SMiKO*^ mice at different time points after TAM induction. As early as 1.0–1.5 months after TAM induction, aortas from *Lkb1*^*SMiKO*^ mice were dramatically dilated, and the medial wall thickness of the aortas was decreased (Fig. 4, A to C). Consistent with scRNA-seq data, VSMCs from *Lkb1*^*SMiKO*^ mice exhibited nearly identical expression of VSMC differentiation markers (α-actin) as WT VSMCs (Fig. 4B). However, VSMCs from *Lkb1*^*SMiKO*^ mice had changed to a flattened, elongated morphology (Fig. 4B) and displayed a corresponding increase in VSMC early-modulation markers and pro-inflammatory genes (fig. S8, A to D), confirming the existence of early-modulated VSMCs during this stage. Moreover, unruffled elastic lamellae were observed during this stage (Fig. 4B).

**Fig.4.**
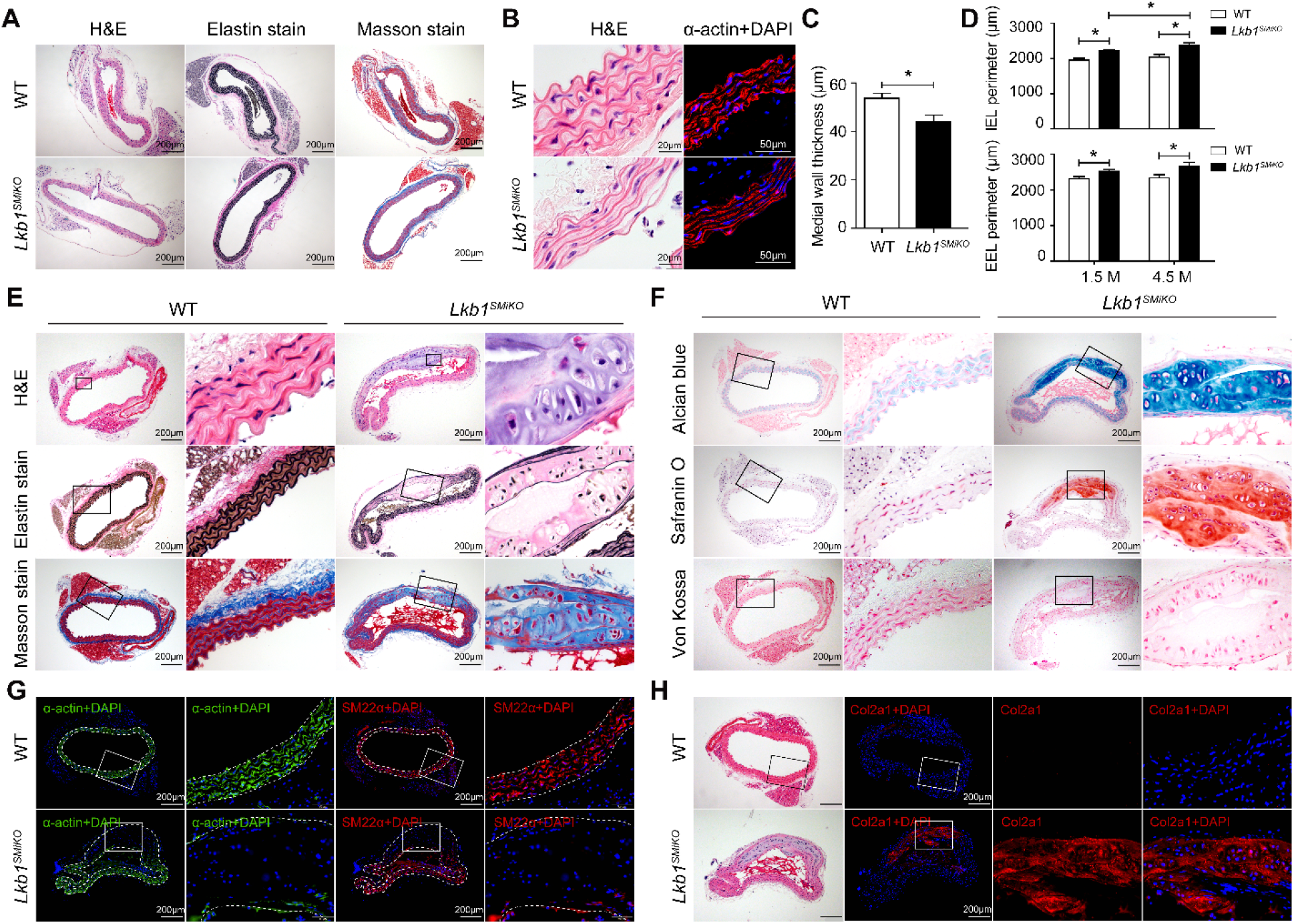
Histological analysis of the aortas from WT and *Lkb1*^*SMiKO*^ mice. **(A)** H&E, elastin, and Masson trichrome stains of the thoracic aortas from WT and *Lkb1*^*SMiKO*^ mice at 1.0–1.5 months post-TAM induction. **(B)** H&E staining and immunofluorescence staining (α-actin) of the thoracic aortas from WT and *Lkb1*^*SMiKO*^ mice at 1.0–1.5 months post-TAM induction. **(C)** Quantification of the medial wall thickness of the thoracic aortas from WT and *Lkb1*^*SMiKO*^ mice at 1.5 months post-TAM induction. **(D)** Quantification of the internal and external elastic laminae (IEL and EEL, respectively) perimeters of the thoracic aortas from WT and *Lkb1*^*SMiKO*^ mice at 1.5 and 4.5 months post-TAM induction. **(E)** H&E, elastin, and Masson stains of the thoracic aortas from WT and *Lkb1*^*SMiKO*^ mice at 4.5 months post-TAM induction. Boxes indicate fields shown at higher magnification in images to the immediate right. **(F)** Alcian blue, Safranin O, and Von Kossa staining of the thoracic aortas from WT and *Lkb1*^*SMiKO*^ mice at 4.5 months post-TAM induction. **(G and H)** Immunofluorescence staining for α-actin (G), SM22α (G), and Col2a1 (H) of the thoracic aortas from WT and *Lkb1*^*SMiKO*^ mice at 4.5 months post-TAM induction. Where indicated, nuclei were counterstained with DAPI. Dashed lines denote the IEL and EEL of the aortas.

We found that the dilation of the aortas from *Lkb1*^*SMiKO*^ mice had progressively increased (Fig. 4D), which was consistent with echocardiography data (Fig. 1, G to J). Notably, there were some unique cells formed in the aorta of *Lkb1*^*SMiKO*^ mice at 4.5 months after TAM induction, which exhibited the typical morphology of chondrocytes (Fig. 4E). In addition, we found that *Lkb1* deletion in VSMCs strongly affected ECM, namely, it led to unruffled elastic lamellae, elastin fiber breakage, and increased collagen deposition (Fig. 4E). The results of the combined Alcian blue-Safranin O staining further revealed the deposition of proteoglycan-rich matrix in *Lkb1*^*SMiKO*^ aorta (Fig. 4F). These data and scRNA-seq data support that, in addition to VSMC transformation, ECM remodeling contributes to the development of vascular disorders in *Lkb1*^*SMiKO*^ mice. Our vascular ultrasound analysis showing that *Lkb1*^*SMiKO*^ mice had increased aortic stiffness (fig. S9) further supported ECM remodeling in *Lkb1*^*SMiKO*^ aorta. However, the Von Kossa staining showed that during this stage (at 4.5 months post-TAM induction) there was no calcification in the thoracic aorta of *Lkb1*^*SMiKO*^ mice (Fig. 4F), suggesting uncalcified cartilaginous metaplasia. As expected, the uniquely transformed cells were characterized by the lack of expression of VSMC markers α-actin and SM-22α (Fig. 4G) and high-level expression of chondrocyte markers Col2a1 (Fig. 4H) and Sox9 (data not shown), consistent with the scRNA-seq data.

When we followed up mice at 6 months post-TAM induction, we found that the aortic structural alterations in *Lkb1*^*SMiKO*^ mice had become more pronounced. The deposition of collagen-rich and proteoglycan-rich matrix extended throughout the entire aortic section (fig. S10, A and B). In addition, there was calcification in the thoracic aorta at this stage, suggesting calcified cartilaginous metaplasia (fig. S10, A and B). Immunofluorescence staining showed that there was no expression of VSMC contraction markers (α-actin and SM-22α) anywhere in the entire aorta section (fig. S10C). By contrast, the expression of chondrocyte or fibroblast markers, such as Col2a1, osteopontin (Opn), and lumican (Lum) was markedly increased in *Lkb1*^*SMiKO*^ aorta (fig. S10D and fig. S11), suggesting that the transformation of VSMCs in *Lkb1*^*SMiKO*^ mice progressively increased.

We further checked other parts of the aorta, including the aortic root and arch. The aortic root of *Lkb1*^*SMiKO*^ mice exhibited uncalcified cartilaginous metaplasia at 4.5 months post-TAM induction (fig. S12A). However, it progressed to calcified cartilaginous metaplasia at 6.0 months post-TAM induction (fig. S12, B and C). For the aortic arch, we found that even at 4.5 months post-TAM induction, calcified cartilaginous metaplasia had formed (fig. S13). Together, these data suggest that different sections of the *Lkb1*^*SMiKO*^ aorta might have different rates and patterns of VSMC transformation progression.

Because *Lkb1* deletion occurs in all vessels that contain VSMCs, we sought to determine the effect of *Lkb1* on some medium-sized arteries (muscular arteries). As early as 1.0–1.5 months after TAM induction, the femoral arteries were dramatically dilated (fig. S14A). With the passage of time, non-ruptured and ruptured femoral artery aneurysms were found in *Lkb1*^*SMiKO*^ mice (fig. S14B). Most importantly, we found that the VSMCs were transformed into fibroblast-like cells that were Dpt-positive and Fbln1-positive (fig. S14C). A similar phenomenon was observed in the renal arteries (data not shown). Therefore, we conclude that loss of *Lkb1* results in the transformation of VSMCs to other cell types, such as fibroblast-like or chondrocyte-like cells. However, different types of arteries or different sections of the artery or aorta might have distinct types of cell transformation.

### Lineage tracing of transformed cell populations in *Lkb1*^*SMiKO*^ mice

To demonstrate the direct transformation of VSMCs into fibroblast-like and chondrocyte-like cells in *Lkb1*^*SMiKO*^ mice, we induced SMC lineage tracing by administrating TAM to triple-transgenic *Lkb1*^*flox/flox*^;*Myh11-Cre/ERT2;ROSA*^*mT/mG*^ mice (fig. S15A). *Lkb1*^*+/+*^;*Myh11-Cre/ERT2;ROSA*^*mT/mG*^ mice were used as controls. tdTomato signals in the vessels were mainly found in the endothelial cells, whereas GFP signals were found in the VSMCs of different kinds of arteries, including aorta, aortic root, femoral artery, renal artery, and coronary artery (fig. S15B). Note that GFP^+^ VSMCs exhibited normal expression of VSMC differentiation marker α-actin throughout the entire thoracic aorta of control mice (Fig. 5A). However, immunostaining revealed a progressive decrease of α-actin expression in GFP^+^ VSMCs in the thoracic aorta of *Lkb1*^*flox/flox*^;*Myh11-Cre/ERT2;ROSA*^*mT/mG*^ mice compared with control mice, which is accompanied by a progressive increase of the proteoglycan-rich matrix deposition (Fig. 5A). More importantly, in thoracic aorta, abdominal aorta, and aortic arch, medial GFP^+^ cells that did not express VSMC differentiation marker α-actin had acquired characteristics of chondrocytes (Fig. 5, A and B, and fig. S16). These data strongly suggest that *Lkb1*^*SMiKO*^ VSMCs directly and gradually transformed into chondrocyte-like cells.

**Fig. 5.**
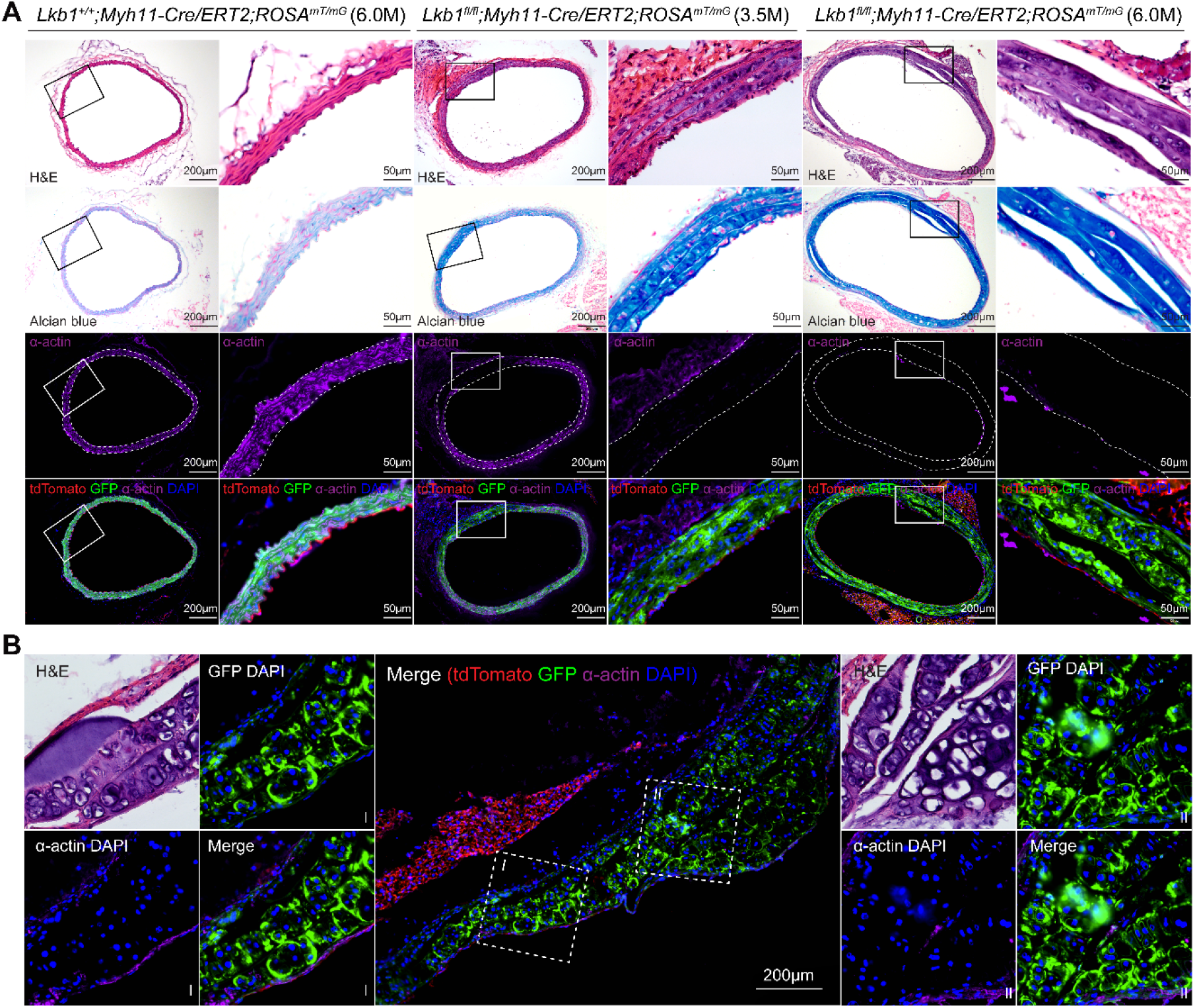
Lineage tracing of VSMCs in *Lkb1*^*SMiKO*^ mice. **(A)** H&E staining, Alcian blue staining, and anti-α-actin co-immunostaining (purple) of thoracic aortas from *Lkb1*^*+/+*^;*Myh11-Cre/ERT2*;*ROSA*^*mT/*^*mG* and *Lkb1*^*fl/fl*^;*Myh11-Cre/ERT2*;*ROSA*^*mT/mG*^ mice at 3.5 and 6.0 months post-TAM induction. Boxes indicate fields shown at higher magnification in images to the immediate right. Dashed lines denote the IEL and EEL of the aortas. **(B)** H&E staining and anti-α-actin co-immunostaining (purple) of aortic arch from *Lkb1*^*fl/fl*^;*Myh11-Cre/ERT2*;*ROSA*^*mT/mG*^ mice at 3.5 months post-TAM induction.

In addition, we also found that in *Lkb1*^*flox/flox*^;*Myh11-Cre/ERT2;ROSA*^*mT/mG*^ mice that developed femoral and renal artery aneurysms, the GFP^+^ cells were characterized by lack of expression of VSMC marker α-actin (fig. S17). Therefore, these data highlighted the important role of progressive VSMC transformation in the pathogenesis of arterial aneurysm.

### Impaired vascular function in *Lkb1*^*SMiKO*^ mice

Given the marked effect of Lkb1 on mouse VSMC fate and vessel morphology changes, we sought to examine the role of Lkb1 in VSMC and arterial function. We first performed blood pressure measurement in WT and *Lkb1*^*SMiKO*^ mice and found that *Lkb1*^*SMiKO*^ mice displayed low blood pressure compared to WT control mice, including systolic, diastolic, and mean blood pressure (Fig. 6A). *Lkb1*^*SMiKO*^ mice also showed dramatically decreased vascular smooth muscle contractility induced by U46619 and phenylephrine (PE), as assessed by myographic testing of isolated mesenteric arterial rings (Fig. 6, B to D).

**Fig. 6.**
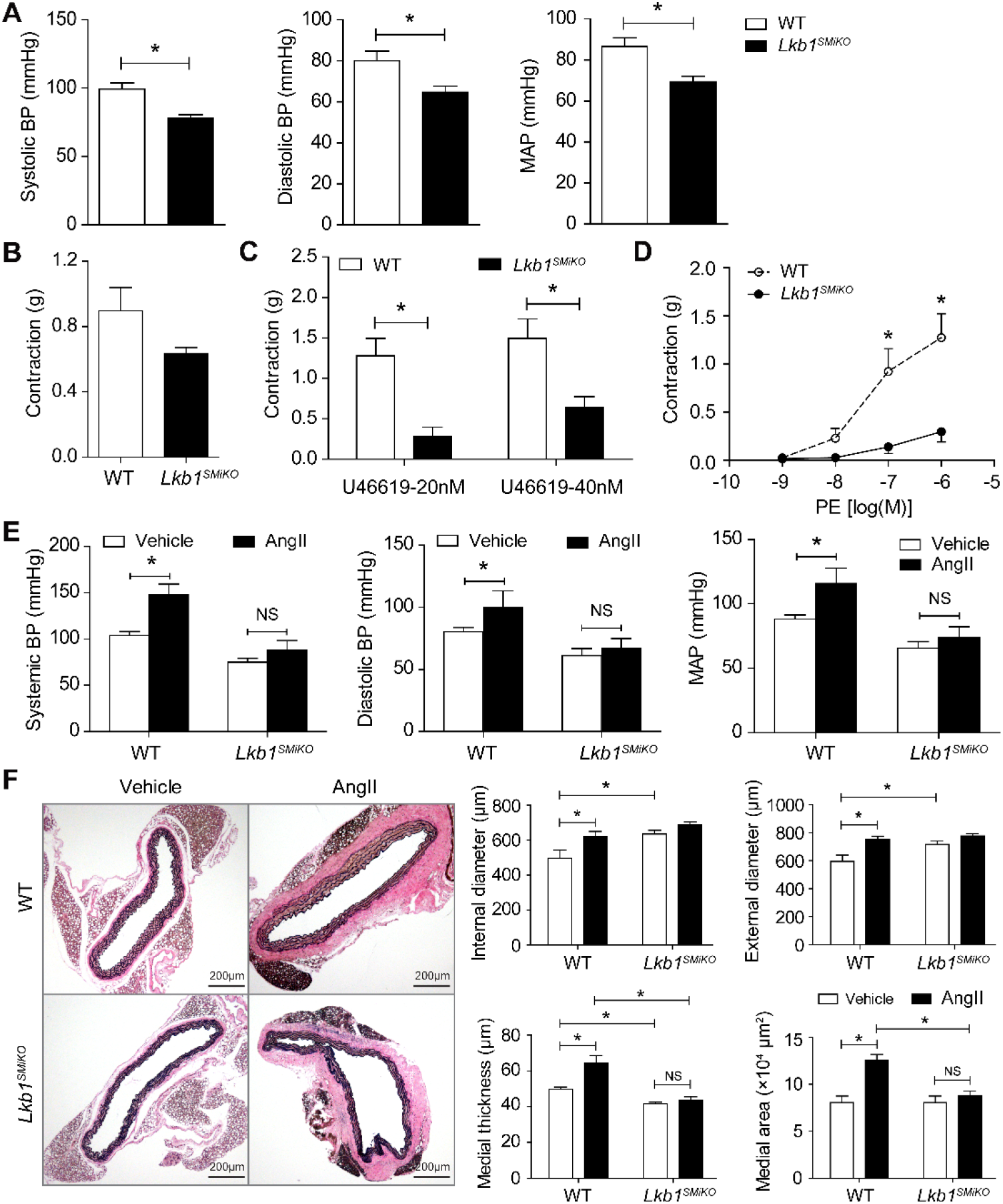
Impaired vascular function in *Lkb1*^*SMiKO*^ mice. **(A)** Blood pressure in WT and *Lkb1*^*SMiKO*^ mice (n=4/group). Blood pressure was measured by inserting fluid-filled catheters into carotid artery. *p<0.05. **(B–D)** The first-order branches of the superior mesenteric arteries were isolated from WT and *Lkb1*^*SMiKO*^ mice (two arteries for each mouse and five mice for each group). Vessel contraction induced by high (60 nM) potassium (B), U46619 (C), and phenylephrine (PE) (D) in control and KO mesenteric arteries was measured using wire myography. **(E)** Arterial blood pressure was measured by inserting fluid-filled catheters into the carotid artery in WT and *Lkb1*^*SMiKO*^ mice infused with vehicle or angiotensin (Ang)II for 4 weeks. **(F)** Representative images of elastin staining and quantification of internal and external diameters, medial wall thickness, and medial area of the thoracic aortas from WT and *Lkb1*^*SMiKO*^ mice infused with vehicle or AngII for 4 weeks.

We further tested the effect of VSMC-specific *Lkb1* ablation on angiotensin II (AngII)-induced hypertension. AngII treatment induced a strong increase in blood pressure in WT control mice, whereas this effect was much more limited in *Lkb1*^*SMiKO*^ mice (Fig. 6E). Interestingly, *Lkb1*^*SMiKO*^ mice had larger internal aortic diameters than those of WT control mice with or without AngII treatment (Fig. 6F). However, VSMC-specific *Lkb1* ablation abolished Ang II-induced aortic medial thickening (Fig. 6F). In addition, AngII-induced cardiac hypertrophy was also less severe in *Lkb1*^*SMiKO*^ mice than in WT control mice (fig. S18). Together, these data suggest that *Lkb1*-deficient VSMCs exhibited a defective response to AngII *in vivo*.

We further checked the role of Lkb1 in the response to injury of the blood vessel wall, employing the left common carotid artery-ligation mouse model. WT arteries showed obvious neointima formation and wall thickening after carotid ligation. Remarkably, the *Lkb1*^*SMiKO*^ arteries developed a severe, organized thrombus that occupied an abnormally enlarged lumen (fig. S19, A and B). Elastic laminae in *Lkb1*^*SMiKO*^ vessels were distended, as indicated by the absence of undulations, and frequently disrupted (fig. S19B). Comparing the medial wall thickness between WT and *Lkb1*^*SMCiKO*^ mice revealed that the media was extremely thin in *Lkb1*^*SMiKO*^ vessels after injury (fig. S19, A to C). Immunofluorescence staining showed the lack of expression of VSMC contraction markers (α-actin) throughout the whole *Lkb1*^*SMiKO*^ vessel section after injury (fig. S19D). These data indicate that *Lkb1*^*SMCiKO*^ vessels were unable to undergo injury-induced neointima formation.

Taken together, these data suggest that Lkb1 deficiency in VSMCs results in hypotension, decreased vascular contractility, and impaired responses to AngII and vessel injury in mice *in vivo*. Because mice used for AngII infusion and carotid artery ligation experiments were 2 weeks post-TAM induction, these data further established that transformed VSMCs, even early modulated VSMCs, would lose the physiological and pathological functions of VSMCs.

## Discussion

Although two decades have passed since the discovery that *Lkb1*^*-/-*^ mice die at mid-gestation with vascular abnormalities, mesenchymal cell death, and neural tube defects (*24*), the role of Lkb1 in VSMC fate and vascular homeostasis remains largely unknown. In this study, we generated two VSMC-specific (*Tagln-Cre* and *Myh11-Cre/ERT2* transgenic) *Lkb1*-deficient mouse models and found that Lkb1 is required for acquisition and maintenance of VSMC fate. We demonstrated that: (1) VSMC-specific *Lkb1* deletion by SM22α-cre leads to severe vascular abnormalities and embryonic lethality. (2) After targeted *Lkb1* ablation by TAM-inducible *Myh11-Cre/ERT2*, mature VSMCs undergo transcriptional reprogramming and transform gradually from early modulated VSMCs to fibroblast-like, chondrocyte-like (cartilage), and even osteocyte-like (calcification) cells, leading to aortic/arterial dilation, aneurysm, and rupture.

Because the phenomenon of VSMC transdifferentiation/transformation has been primarily studied by stimulating cultured VSMCs with lipids and various growth factors (*28–30*) or using mouse models of atherosclerosis (*8, 31–34*), here we provide a unique *in vivo* mouse model of spontaneous VSMC transformation. Most importantly, by combining scRNA-seq, lineage tracing, and histological analysis, we clearly displayed several distinct transformed cell types or states that occur at different stages of VSMC transformation. We found that VSMCs that undergo cellular transformation appear to exhibit a continuous trajectory from early modulated VSMCs to fibroblast-like and chondrocyte-like cells, which is driven by a highly dynamic transcriptional reprogramming. In particular, we defined the early modulated VSMC as an intermediate cell state during VSMC transformation for the first time. Moreover, we found that the *Lkb1* deletion-induced transformed VSMCs would lose the physiological and pathological functions of VSMCs, even in early stage as early modulated VSMCs. Therefore, our work provides a valuable mouse model to investigate the continuous and progressive process of VSMC transformation.

We also found that *Lkb1*^*SMiKO*^ mice spontaneously developed arterial aneurysms in abdominal aorta, renal artery, iliac artery, femoral artery, and/or popliteal artery, which is highly reminiscent of human aneurysms (*35*). Therefore, we also provide an ideal animal model to capture the trajectory of arterial aneurysm formation that resembles human anatomy and pathophysiology. Moreover, our results shed new light on the role of VSMC transformation in arterial aneurysm and suggest that progressive VSMC transformation is the driving force for arterial dilation and aneurysm formation.

Our findings suggest that, in addition to VSMC transformation, ECM remodeling plays important roles in vascular disorders. We found that Lkb1 deficiency in VSMCs led to striking ECM remodeling. In *Lkb1*^*SMiKO*^ aorta, the elastic lamellae become unruffled, fragmented, and discontinuous. Most importantly, *Lkb1*^*SMiKO*^ aorta exhibited excessive deposition of collagen-rich and proteoglycan-rich matrix, accompanied by increased aortic stiffness. These observations provide clear evidence supporting the notion that segmental aortic stiffening is an early pathomechanism for aneurysm progression (*36*). Further, we also provided data showing the increase in inflammatory cell infiltration during this process. Taking these data together, we speculate that VSMC transformation promotes ECM remodeling and inflammatory cell infiltration, which may further accelerate the transformation of VSMCs, leading to aneurysm formation.

In conclusion, the present study introduces the novel concept that VSMC transformation followed by ECM remodeling and inflammatory cell infiltration contributes to aortic or arterial aneurysm formation, and that Lkb1 is the predominant regulator of mouse VSMC fate maintenance. In other words, aneurysm progression can be regarded as a complex biological process involving VSMC-transformation–driven remodeling of the vessel wall. An improved understanding of VSMC fate regulation may ultimately lead to new approaches to prevent and treat arterial dilation and aneurysm.

## Supporting information

Cai-Supplement

## Acknowledgments

This study was supported by the National Institutes of Health grants (HL079584, HL080499, HL089920, HL110488, HL128014, HL132500, HL137371, HL140954, HL142287, AG047776, and CA213022). This work was, in part, supported by the Georgia Research Alliance. M.-H.Z. is a Georgia Research Alliance Eminent Scholar in Molecular Medicine.

## Author contributions

Z.C. designed and executed the experiments, analyzed the data and wrote the manuscript. P.S., S.Y., and Z.L. provided advice and analyzed the data. F.Z., J.M., X.Z., Y.D., L.X., T.R., and Y.Q. provided technical assistance and advice. B. H. provided advice, analyzed the data and revised the manuscript. M.-H.Z. conceived and designed the experiments, analyzed the data, and revised the manuscript.

## Supplementary Materials

Materials and Methods

Figures S1-S19

Tables S1-S2

References (*37-47*)

