## Supplementary material for "Lkb1 is a master regulator of VSMC fate and function in mice": Cai-Supplement

**This PDF file includes:**

Materials and Methods

Figs. S1 to S19

Tables S1 to S2

### Materials and Methods

Mouse lines

*Tagln-Cre* transgenic mice (Stock No. 017491), *Myh11-Cre/ERT2* transgenic mice (Stock No. 019079) and Lkb1-floxed (*Lkb1$^{flox/flox}$*) mice (Stock No. 014143) were purchased from The Jackson Laboratory. *Tagln-Cre* mice and *Myh11-Cre/ERT2* mice were intercrossed with *Lkb1$^{flox/flox}$* mice to generate *Lkb1$^{flox/flox}$;Tagln-Cre* and *Lkb1$^{flox/flox}$;Myh11-Cre/ERT2* mice, respectively. *Lkb1$^{flox/flox}$;Myh11-Cre/ERT2* mice were further bred with *ROSA$^{mT/mG}$* mice (The Jackson Laboratory, Stock No. 007676) to generate triple-transgenic *Lkb1$^{flox/flox}$;Myh11-Cre/ERT2;ROSA$^{mT/mG}$* mice. *Lkb1$^{flox/flox}$;Myh11-Cre/ERT2* and *Lkb1$^{flox/flox}$;Myh11-Cre/ERT2;ROSA$^{mT/mG}$* mice aged 4–8 weeks were intraperitoneally injected with 100 µl tamoxifen solution (1 mg in 100 µL sunflower oil per mouse) for 5 consecutive days to specifically activate Cre recombinase in mature VSMCs. Mice were housed in a controlled environment (22°C, 12-h/12-h light/dark cycle) with free access to food and water. All mouse experimental protocols were approved by the Institutional Animal Care and Use Committee at Georgia State University and were in compliance with relevant ethical regulations.

Animal models

Partial ligation of the left common carotid artery (LCCA) was performed as described previously (*37*). For AngII perfusion, mice were infused with AngII (1,000 ng/kg/min) or physiological saline (0.9% sodium chloride) by Alzet osmotic pumps (DURECT Corp, Model 2004) for 4 weeks as described previously (*38*). Animal models were generated in WT and *Lkb1$^{SMiKO}$* mice at 2 weeks post-TAM induction.

Vascular ultrasound

Vascular ultrasound was performed using a Vevo 3100 Imaging System (FUJIFILM VisualSonics, Toronto, ON, Canada). During the imaging, the mice are anesthetized using isoflurane 1%−3% vol/vol and placed on a heating platform to reduce procedural stress and prevent hypothermia. The ascending aorta and abdominal aorta were visualized in brightness (B)-mode. Aortic wall motion was then recorded in perpendicular orientation using M-mode images captured at specific locations along the infrarenal aorta. Similar to the aorta, the carotid artery was first visualized on the transverse plane in B mode, and then the transducer was switched to obtain images on the longitudinal plane. Anterior and posterior aortic wall motion was assessed using images captured in M mode. Images were analyzed using the Vevo WorkStation. Systolic diameter (Ds) and diastolic diameter (Dd) were quantified from M-mode images, and then circumferential cyclic strain was calculated as (Ds–Dd)/Dd (*36, 39*). The aortic pulse wave velocity (PWV) was calculated as the distance between 2 measurement points divided by the time shift of the waveforms at the 2 points (*40*). For the abdominal aorta, the blood flow and physiological signals were recorded along the abdominal aorta, focusing on 2 points, at the suprarenal branch and ≈1 mm before the left renal artery branch. The time lag in blood flow at these 2 sites was measured between the R waves in electrocardiography (ECG). Peak blood flow ($\Delta t_1$, $\Delta t_2$) time was recorded. $\Delta t_1$ was the time lag in the low site (suprarenal) and $\Delta t_2$ the time lag in the upper site (≈1 mm before the renal artery). The time required for the wave to go from the upper site to the low point provided the pulse transit time (PTT), calculated as follows: $PPT=\Delta T=\Delta t_1-\Delta t_2$. The distance (S) traveled between these two sites was measured using color Doppler images. Pulse wave velocity (PWV) was calculated as follows: $PWV=S/PPT=S/\Delta T=S/(\Delta t_1-\Delta t_2)$. All echocardiography procedures, including data acquisition and analysis, were performed by a researcher blinded to the identity of the samples. Doppler spectrograms of aortic flow at the carotid artery and abdominal aortic site were acquired with a 20-MHz pulsed Doppler probe using the Vevo 3100 Imaging System.

Blood pressure measurement

Blood pressure was measured by inserting fluid-filled catheters into the carotid artery as described previously (*25, 41*). Mice were anesthetized with a mixture of ketamine and xylazine (70:6 mg/kg, intraperitoneal injection) and placed under a heat lamp at 37°C. A catheter was inserted into the left common carotid artery with the aid of a dissecting microscope to measure arterial blood pressure. For catheter insertion, the left common carotid artery was carefully exposed via a 0.5- to 1.0-cm midline incision in the ventral neck region. The tip of the artery toward the head was ligated with a suture (5-0 silk), and the tip toward heart was occluded with a microclip (No. 18055-03; Fine Science Tools, Foster City, CA). A small cut was then made in the vessel wall using microscissors (No. 15000-08, Fine Science Tools). A 60-cm catheter (PE10 tubing; A-M Systems, Sequim, WA) containing a sterile 10% heparin-90% saline solution was inserted into the artery at a distance of 0.65 cm toward the thorax. The arterial clip was removed, and the catheter was tied in place. Blood was directed to a pressure transducer through the catheter to obtain computerized blood pressure measurements (ADInstruments). The mice were allowed to recover, and the systolic and diastolic blood pressures were monitored for at least 30 min in the conscious state.

Measurement of vessel tension in mice

Mice were anesthetized with isoflurane (Covetrus, Dublin, Ohio) and killed by decapitation. Mesenteric arteries were rapidly removed, immersed in Krebs bicarbonate buffer (118 mM NaCl, 4.7 mM KCl, 25 mM $NaHCO_3$, 1.2 mM $KH_2PO_4$, 1.2 mM $MgSO_4$, 2.5 mM $CaCl_2$, and 5 mM glucose), gassed with a mixture of 95% $O_2$ and 5% $CO_2$, and carefully cleaned of all fat and connective tissue. The endothelium was gently removed using a cotton stick. Artery rings were mounted between two hooks in a 5-ml, 37°C organ bath perfused with Krebs buffer. After undergoing an equilibration period, rings were contracted with 60 mM (high) potassium salt solution. The rings were then washed, subjected to an additional equilibration period (30 min), and contracted with U46619 or phenylephrine (PE).

Tissue collection and processing

Tissue sections were stored at -80°C for RNA extraction or western blotting. Tissue sections for pathological diagnosis or immunohistochemistry/immunofluorescence were fixed in 10% neutral buffered formalin and embedded in paraffin or optimal cutting temperature (OCT) compound. Paraffin-embedded sections or OCT-embedded sections were cut at 5 μm or 8 μm thickness, respectively.

Histology and immunohistochemistry

Paraffin-embedded tissue sections were stained with hematoxylin and eosin (H&E) according to standard protocols. All images were recorded using an Olympus digital camera (Tokyo, Japan). For immunohistochemical staining, paraffin-embedded tissue sections were deparaffinized, rehydrated, and subjected to antigen retrieval. Endogenous peroxidase activity was blocked using 0.3% $H_2O_2$ for 20 min. Tissue sections were blocked with normal goat serum (Biogenex, HK112-9K) and incubated with primary antibodies against telethonin (Thermo Fisher, Cat# PA5-78255, 1:100), osteopontin (Abcam, Cat# ab8448, 1:100), lumican (Abcam, Cat# ab168348, 1:100), Dermatopontin (Proteintech, Cat# 10537-1-AP, 1:100), and Fibulin1 (Thermo Fisher, Cat# PA5-103841, 1:200).. Goat anti-rabbit/mouse IgG (DAKO, Cat #K4061, ready-to-use) or goat anti-rat IgG (Millipore, Cat #AP136P, 1:250) were used as secondary antibodies. The reaction was visualized using DAB (DAKO, Cat #K3468) and sections were counterstained with hematoxylin.

Immunofluorescence

Immunofluorescence staining was performed as previously described (*42, 43*). Briefly, OCT-embedded sections were washed with phosphate-buffered saline (PBS), fixed with acetone at 4°C for 15 min, and permeabilized with 0.2% Triton X-100 for 10 min. After blocking with goat serum (protein block) for 30 min, the sections were incubated with primary antibodies against α-actin (Abcam, Cat #ab7817, 1:200), SM22α (Abcam, Cat #ab14106, 1:200), Col2A1 (Boster, Cat #PA2141-1, 1:200), and Sox9 (Millipore, Cat #ABE2868, 1:500) overnight. Secondary antibodies (Alexa Fluor® 488 goat anti-mouse, Alexa Fluor® 555 goat anti-rabbit, and Alexa Flour® 647 goat anti-rat) were added for 1 h at 37°C, followed by nuclear DNA staining using DAPI for 10 min. Fluorescence signals were evaluated using confocal microscopy (LSM 810, Zeiss, Oberkochen, Germany) or immunofluorescence microscopy.

Special stains

Van Gieson Solution Acid Fuchsin (Cat #HT25A; Sigma-Aldrich, St. Louis, MO) was used for elastic fiber staining in mouse aorta samples. Masson trichrome staining (Cat #HT15, Sigma-Aldrich) was performed to measure collagen. Alcian blue and Safranin O staining were performed to detect glycosaminoglycans (GAGs) and chondrogenic differentiation of VSMCs. Briefly, the aorta samples were incubated with 1% Alcian blue at pH 1.0 (Cat #05500, Sigma-Aldrich) followed by nuclear fast red counterstain or 0.1% Safranin O solution (Cat #TMS-009-C, Sigma-Aldrich) followed by Weigert's iron hematoxylin. Von Kossa staining and Alizarin red staining (American MasterTech Scientific, Lodi, CA, USA) were performed for detection of calcium deposits in aorta samples and according to the manufacturer's instructions.

Western blotting

Total protein was prepared from cultured cells, and western blotting was performed as briefly described below. Proteins were quantified with a Pierce BCA Protein Assay Kit, separated by SDS-PAGE, and transferred to nitrocellulose membranes. Membranes were blocked with 5% non-fat milk dissolved in Tris-buffered saline with Tween 20 (TBST) at 37°C for 1 h, and incubated with primary antibodies at 4°C overnight. After incubation with horseradish peroxidase-conjugated secondary antibodies at 37°C for 1 h, proteins were detected using Pierce ECL Western Blotting Substrate and quantified using Quantity One 4.4.0 software (Bio-Rad, Hercules, CA, USA).

RNA extraction and quantitative (real-time) PCR (qPCR)

Total RNA was extracted from mouse tissues using Trizol reagent (Invitrogen), according to the manufacturer's instructions. Total RNA (1 μg) was reverse-transcribed into first-strand cDNA, and qPCR amplification was performed using SYBR Green dye and the CFX96 Touch$^{TM}$ Real-time PCR Detection System (Bio-Rad). The primer sequences used are presented in Table S2. Relative mRNA expression was calculated using the comparative ΔΔCT method and the resulting values were normalized to *18S* ribosomal RNA expression. PCR was performed in triplicate for each experiment. The results presented represent three independent experiments.

Preparation of aorta single-cell suspension

Single-cell suspensions of whole aorta were prepared as previously reported (*44*) with minor modifications. In brief, after mice were perfused with at least 10 mL of fresh cold Hanks' Balanced Salt Solution (HBSS), the whole aorta (including the aortic root, arch, thoracic aorta and abdominal aorta) was carefully dissected. The perivascular adipose tissue was carefully removed under a dissecting microscope. The aortic tissue was cut into 2–5 mm pieces and digested in HBSS containing 2 mg/ml collagenase I, 1 mg/ml collagenase XI, 120 U/ml hyaluronidase, and 100 μg/ml DNase I for 1.5 h at 37°C. The digested aortic suspension was subsequently filtered through a 70-μm cell strainer (Corning) and washed with pre-warmed Dulbecco's Modified Eagle Medium (DMEM)/F-12 50/50 supplemented with 10% fetal calf serum (FCS), 100 U/ml penicillin, and 100 U/ml streptomycin. Cells were centrifuged at 200*g* for 5 min, resuspended in DMEM/F-12 50/50 medium with supplement, and kept in a cell culture incubator for 30 min to allow for recovery. Cells were counted, frozen in standard freezing media (90% fetal bovine serum [FBS]+10% DMSO), and stored at -80°C using a cell-freezing container, until used for single-cell library preparation.

Single-cell capture and library preparation

Single-cell (sc) RNA-seq was performed by microfluidic inDrop encapsulation, barcoding, and library preparation, as previously described (*45, 46*). The scRNA-Seq libraries were generated from each experimental time point using the 10x Genomics Chromium Controller and Chromium Single Cell 3' V2 (samples from WT-1M and KO-1M) and 5'PE (samples from WT, KO-3.5M, and KO-4.5M) Reagent Kits (10X Genomics, Pleasanton, CA) per the manufacturer's instructions. Reverse transcription and sample indexing were performed using the C1000 Touch Thermal cycler with 96-Deep Well Reaction Module (Thermo Fisher Scientific, Waltham, MA). Briefly, the suspended cells were concentrated to a density of 1000 cells/μL and approximately 10,000 single cells were loaded into each channel to first generate single-cell gel beads in emulsion (GEMs). After breaking the GEMs, the barcoded cDNA was purified and amplified. The amplified barcoded cDNA was fragmented, A-tailed, and ligated to adaptors. Finally, PCR amplification was performed to enable sample indexing and enrichment of the RNA-seq libraries. The final libraries were quantified using the Qubit High Sensitivity DNA assay (Thermo Fisher Scientific) and the size distribution of the libraries was determined using a High Sensitivity DNA chip on a Bioanalyzer 2200 (Agilent, Santa Clara, CA). All libraries were sequenced on an Illumina HiSeq 4000 instrument (Illumina, San Diego, CA).

Analysis of scRNA-Seq data

ScRNA-seq data analysis was performed by NovelBio (Shanghai, China) with the NovelBrain Cloud Analysis Platform. We applied fastp (*47*) with default parameter filtering the adaptor sequence and removed the low-quality reads to achieve clean data. For the 10x data, the feature-barcode matrices were obtained by aligning reads to the mouse genome (Ensembl version 92) using Cell Ranger v3.0.0 (10x Genomics). The Seurat package (version: 3.1.4, https://satijalab.org/seurat/) was applied for cell normalization and regression based on the expression table according to the unique molecular identifier (UMI) counts of each sample and percent of mitochondria to obtain the scaled data. Cells containing greater than 200 expressed genes and mitochondria UMI rate less than 20% passed the cell-quality filtering and mitochondrial genes were removed from the expression table. Principle component analysis (PCA) was constructed with the top 2000 variable genes and the top 10 principals were used for a construction. Utilizing graph-based clustering, we acquired the unsupervised cell cluster result based on the PCA top 10 principals and we calculated the marker genes using the FindAllMarkers function with the Wilcox rank sum test algorithm and the following criteria: 1) lnFC > 0.25; 2) P value<0.05; 3) min.pct>0.1. In particular, to integrate cells from different reagent kits for unsupervised clustering, robust PCA (RPCA) was used in the Seurat alignment workflow (version: 3.1.4, https://satijalab.org/seurat/). In order to identify the cell type detailed, clusters of the same cell type were selected for re-tSNE analysis, graph-based clustering, and marker analysis.

Pseudotime Analysis

We applied single-cell trajectory analysis using Monocle 2 (http://cole-trapnell-lab.github.io/monocle-release) using DDR-Tree and default parameters. Before Monocle analysis, we selected marker genes of the Seurat clustering result and raw expression counts after cell filtering. Based on the pseudotime analysis, branch expression analysis modeling (BEAM analysis) was applied for branch-fate–determined gene analysis.

Quantification and statistical analysis

Statistical analyses were performed using GraphPad Prism 5.0 software (GraphPad, San Diego, CA). Unpaired two-tailed Student's *t*-tests were used to calculate significant differences between two groups. Multiple comparison correction analysis was performed using one-way ANOVA with Tukey's post hoc HSD test. A log-rank (Mantel-Cox) test was used to analyze Kaplan–Meier curves. Fisher's exact test was applied to test for association between categorical variables. $P < 0.05$ was considered statistically significant.

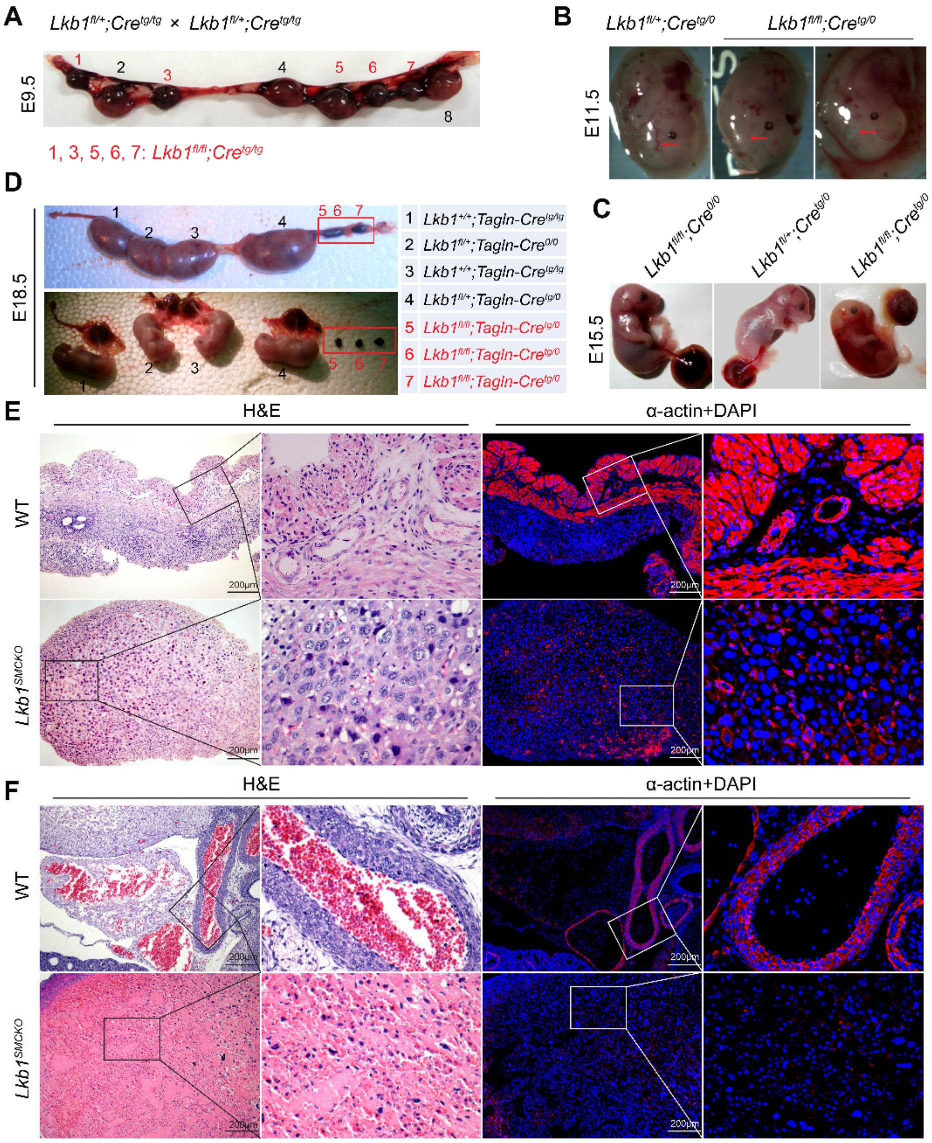


**Figure S1. Embryos from *Lkb1*$^{SMKO}$ mice showed severe vascular abnormalities**

(A–D) Representative images of *Lkb1*$^{flox/flox}$;*Tagln-Cre*, *Lkb1*$^{flox/+}$;*Tagln-Cre* and wild type (WT) control embryos at different embryonic days (E9.5, E11.5, E15.5, and E18.5).

(E and F) Hematoxylin-and-eosin (H&E) and anti–α-actin immunofluorescence (IF) staining of transverse sections of WT and *Lkb1*$^{SMKO}$ embryos at embryonic day (E)8.5 and 14.5 (E8.5 and E14.5).

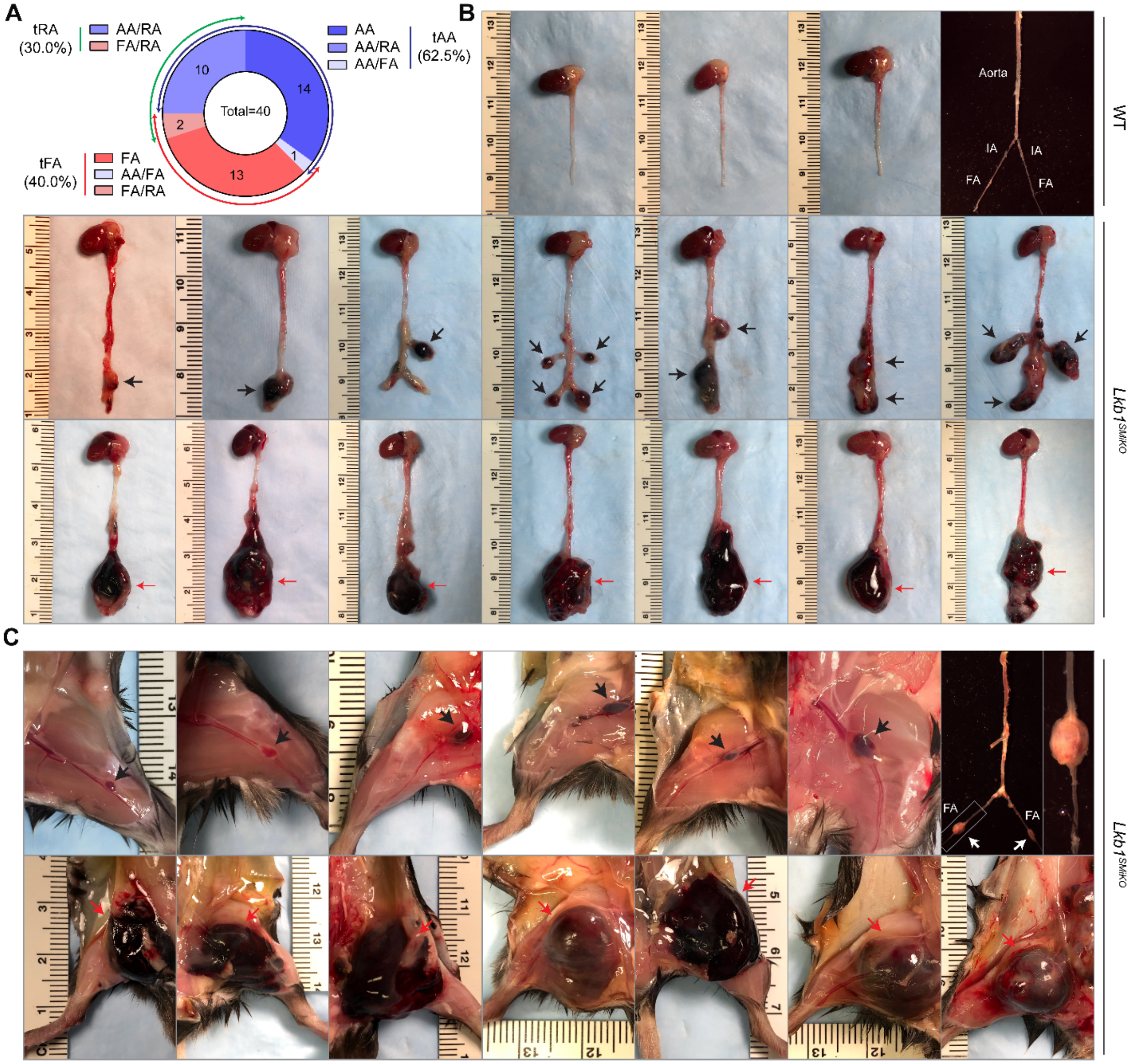


**Figure S2.** ***Lkb1***$^{SMiKO}$ **mice developed aortic and/or arterial aneurysm and rupture**

(A) Pie chart showing the incidence of aortic and/or arterial rupture in *Lkb1*$^{SMiKO}$ mice. AA, abdominal aortic rupture; RA, renal arterial rupture; FA, femoral and/or popliteal arterial rupture; AA/RA, ruptures of the abdominal aorta and renal artery; AA/FA, ruptures of the abdominal aorta and femoral/popliteal artery; FA/RA, ruptures of the femoral/popliteal artery and renal artery.

(B and C) Representative images of aneurysms and ruptures in abdominal aorta (B) and femoral/popliteal artery (C) in *Lkb1*$^{SMiKO}$ mice, along with normal aorta from WT mice. Black or white arrows denote aneurysm. Red arrows denote the ruptured aorta/artery. FA, femoral artery; IA, iliac artery.

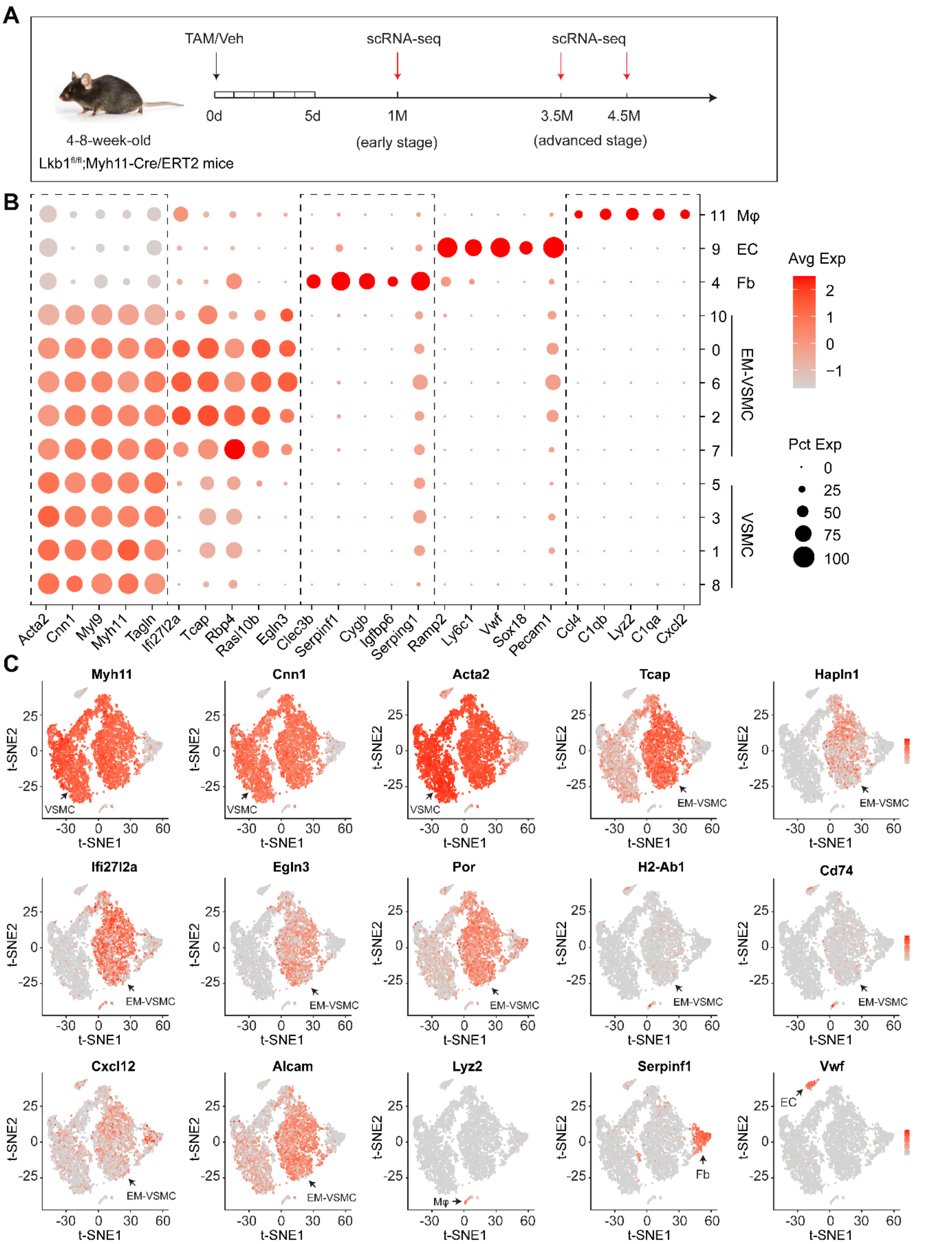


**Figure S3. Gene expression differences between the early-stage aortas of WT and *Lkb1*$^{SMiKO}$ mice**

(A) Design of single-cell RNA sequencing (scRNA-seq) experiment.

(B) Bubble plot of gene expression of five representative marker genes in single aortic cells isolated from WT and *Lkb1*$^{SMiKO}$ mice at 1.0 month (WT-1M and KO-1M, respectively) post-TAM induction. Dot color scale indicates the average expression level (Avg Exp) and dot size represents percentage of cells in each cluster that express at least one transcript of each gene (Pct Exp).

(C) *t*-SNE plot showing expression levels of the indicated genes. Black arrows indicate cells with high expression of the indicated marker gene.

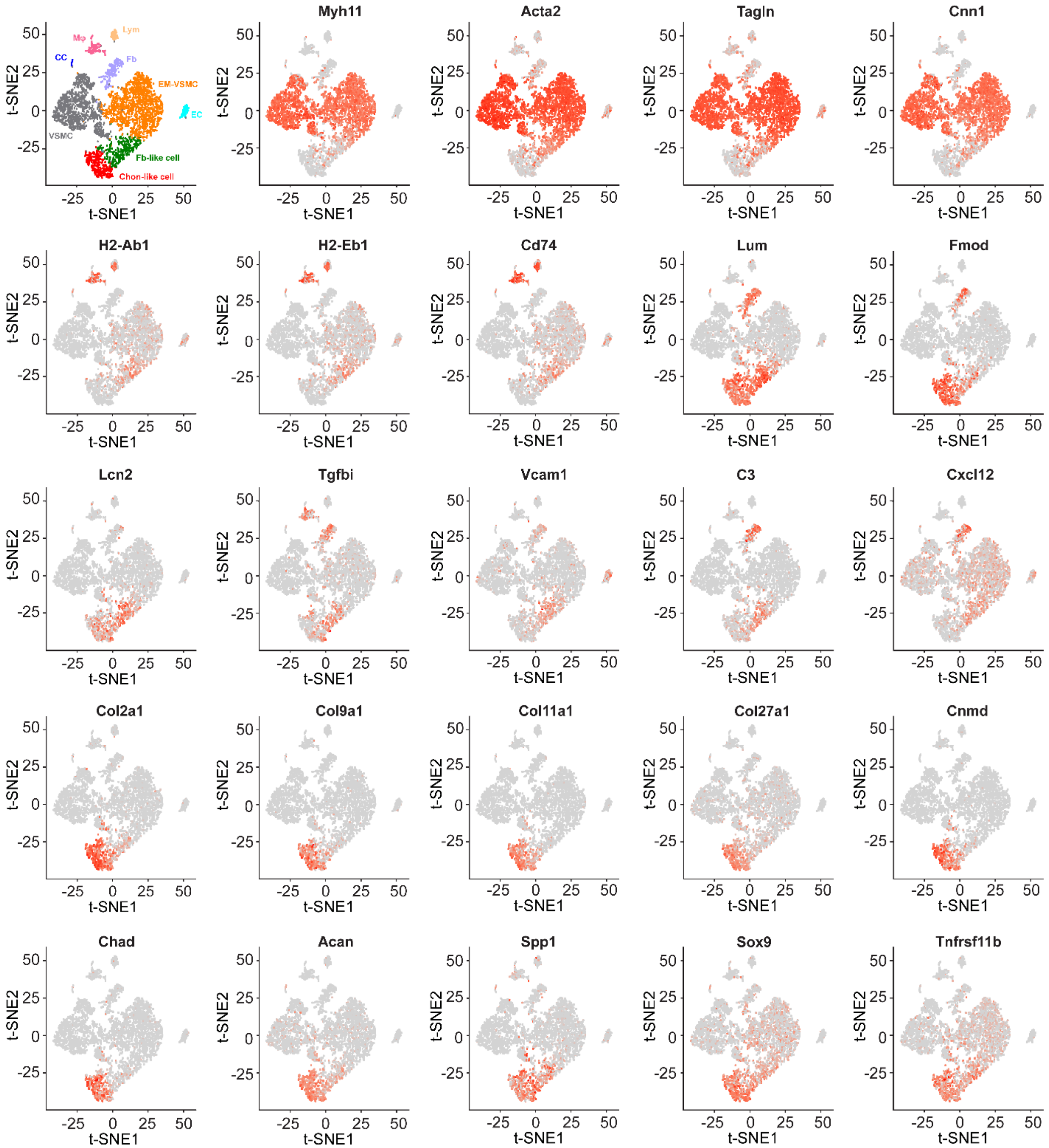


**Figure S4. Gene expression differences between the advanced-stage aortas of WT and *Lkb1*$^{SMiKO}$ mice**

*t*-SNE plot showing expression level of the indicated genes in single aortic cells isolated from WT and *Lkb1*$^{SMiKO}$ mice at 3.5 and 4.5 months (WT-4.5M, KO-3.5M, and KO-4.5M, respectively) post-TAM induction.

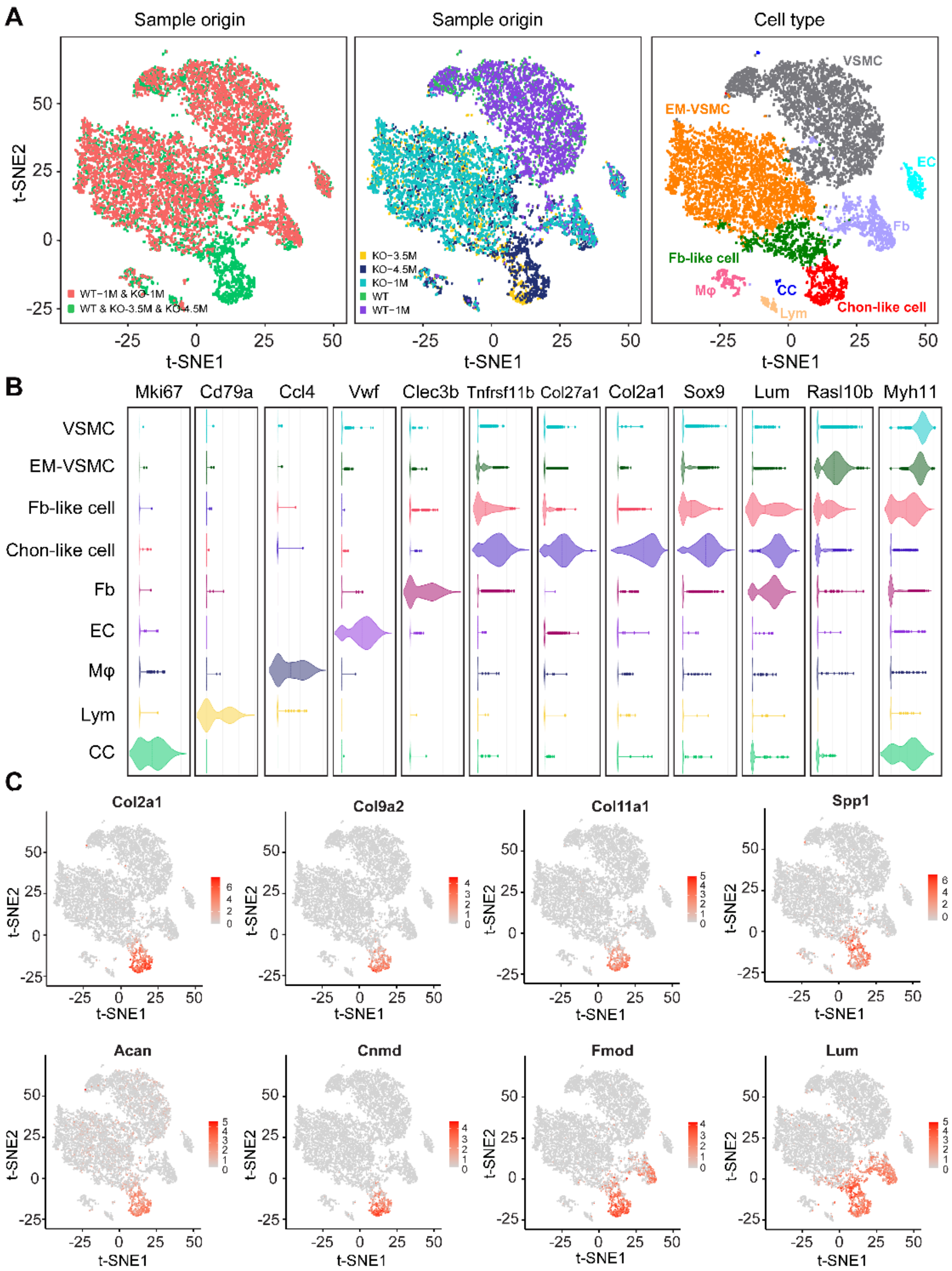


**Figure S5. Joint clustering of the early- and advanced-stage aortas of WT and *Lkb1*$^{SMiKO}$ mice**

(A) Joint clustering of early- and advanced-stage aortas of WT and *Lkb1*$^{SMiKO}$ mice using robust principal component analysis (RPCA) as per the Seurat R package, colored by sample origin and cell type (VSMC, vascular smooth muscle cells; EM-VSMC, early modulated vascular smooth muscle cells; EC, endothelial cell; Fb, fibroblast; Mφ, macrophage; Lym, lymphocyte; CC, cycling cell; Fb-like cell, fibroblast-like cell; Chon-like cell, chondrocyte-like cell).

(B) Violin plots showing expression levels of canonical marker genes.

(C) *t*-SNE plot showing expression level of the indicated genes.

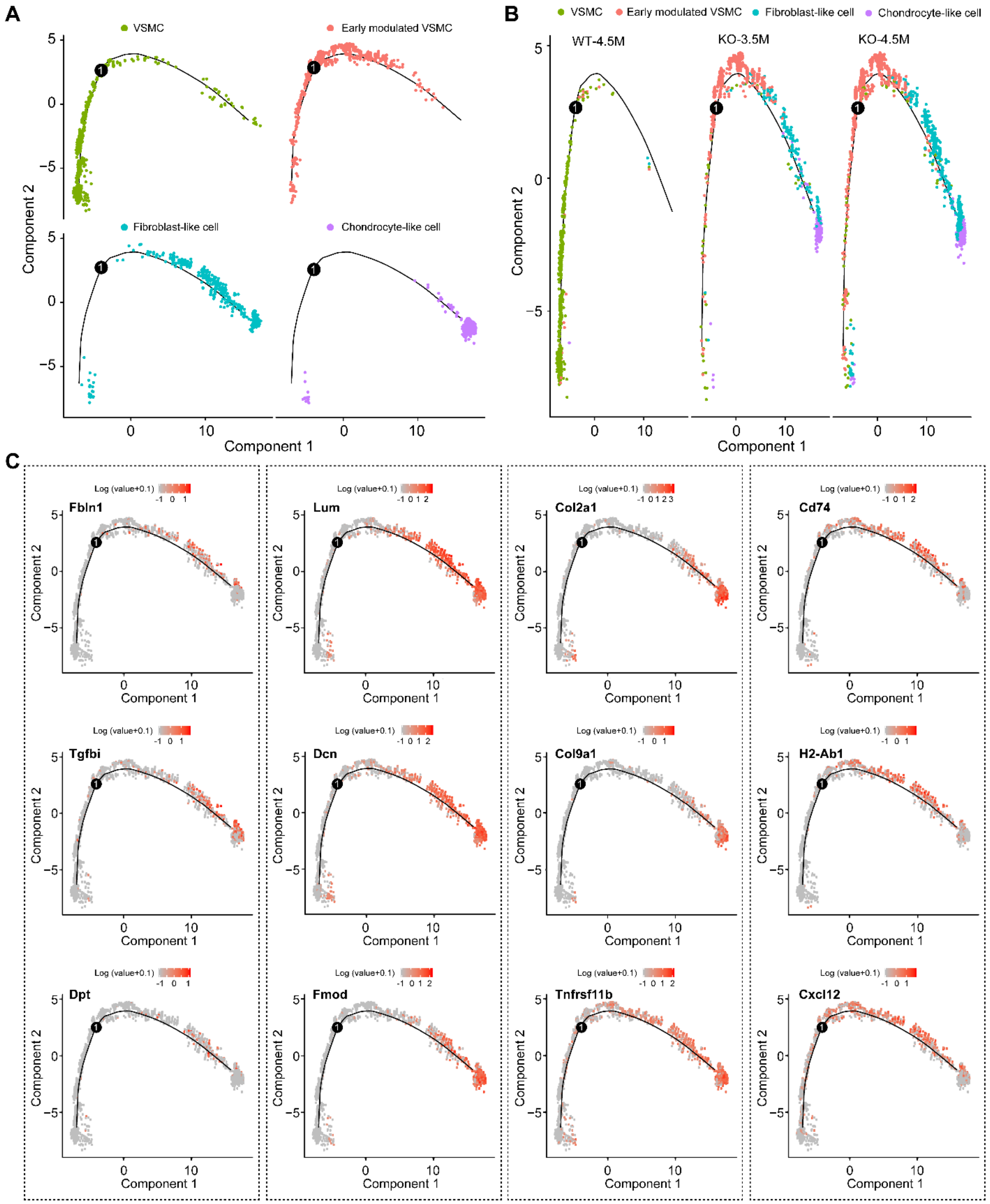


**Figure S6. Single-cell trajectories reveal a gradual VSMC transformation.**

(A-B) Monocle 2 pseudotime analysis of VSMCs, early modulated VSMCs (EM-VSMC), fibroblast-like, and chondrocyte-like cells.

(C) Expression levels of the indicated genes along the pseudotime trajectory.

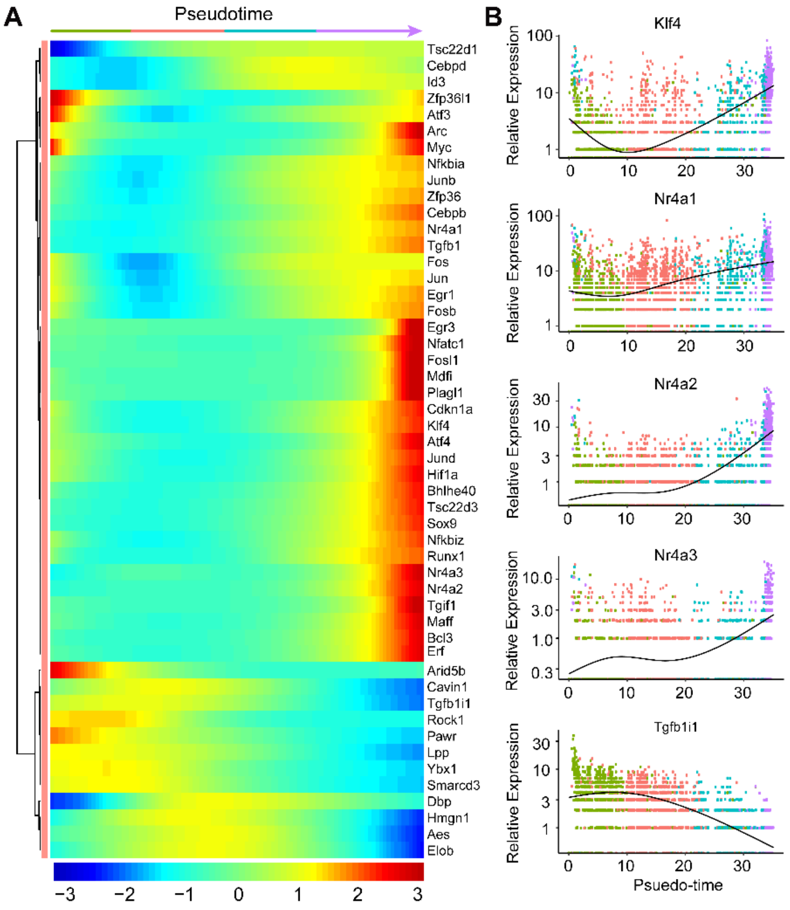


**Figure S7. Single-cell trajectories reveal dynamic transcriptional reprogramming**

(A) Heatmap showing top 50 differentially expressed transcription factors, ordered based on their common kinetics. The Expression Z score indicates changes in a gene relative to its dynamic range over pseudotime.

(B) Kinetic diagrams showing the expression of some genes over pseudotime.

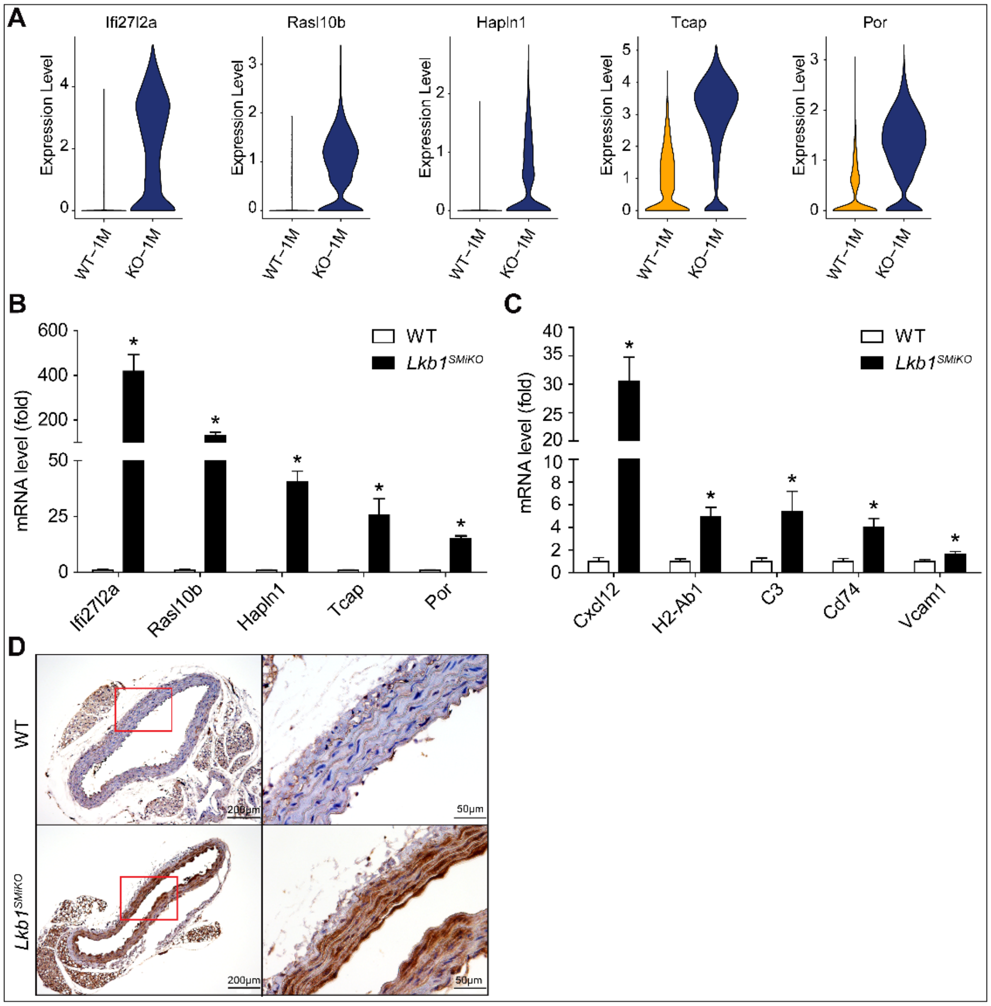


**Figure S8. VSMC early-modulation–associated genes were increased in the early-stage aortas of *Lkb1*$^{SMiKO}$ mice**

(A) Violin plots showing expression levels of early-modulation–associated genes from single aortic cells isolated from WT and *Lkb1*$^{SMiKO}$ mice at 1.0 month (WT-1M and KO-1M) post-TAM induction.

(B and C) Real-time PCR analysis of the indicated genes from the aortas of WT and *Lkb1*$^{SMiKO}$ mice at 1.0–1.5 months post-TAM induction.

(D) Immunohistochemical analysis of Tcap (telethonin) in the thoracic aortas of WT and *Lkb1*$^{SMiKO}$ mice at 1.0–1.5 months post-TAM induction.

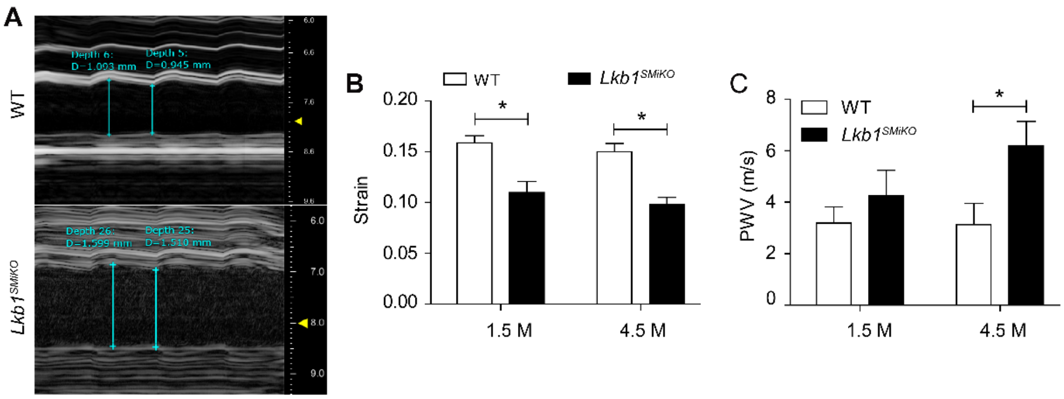


**Figure S9.** ***Lkb1$^{SMiKO}$*** **mice had increased aortic stiffness.**

(A) Representative images of motion (M)-mode for the aorta monitored by ultrasound.

(B and C) Circumferential cyclic strain (B) and analysis of segmental stiffness (C) of the abdominal aorta in WT and *Lkb1$^{SMiKO}$* mice (n=8–10).

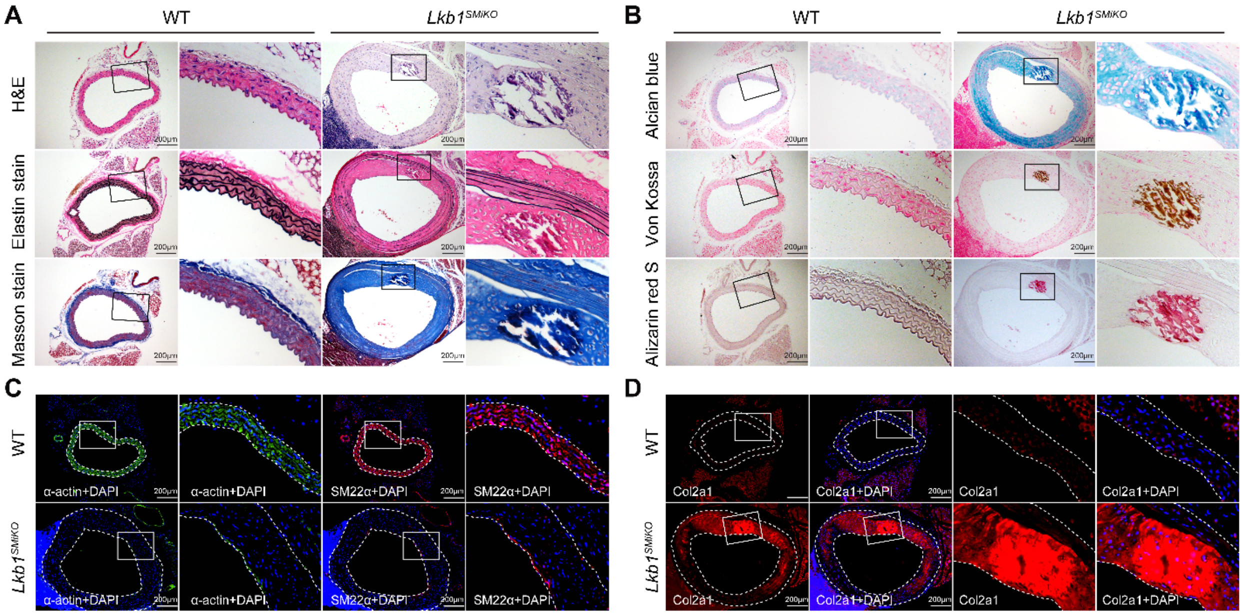


**Figure S10. Histological analysis of the aortas of WT and *Lkb1*$^{SMiKO}$ mice.**

(A) H&E, elastin, and Masson stains of the thoracic aortas of WT and *Lkb1*$^{SMiKO}$ mice at 6 months post-TAM induction.

(B) Alcian blue, Von Kossa, and Alizarin red S stains of the thoracic aortas of WT and *Lkb1*$^{SMiKO}$ mice at 6 months post-TAM induction.

(C and D) IF staining for α-actin (C), SM22α (C), and Col2a1 (D) in the thoracic aortas of WT and *Lkb1*$^{SMiKO}$ mice at 6 months post-TAM induction. Dashed lines denote the internal and external elastic laminae of the aortas.

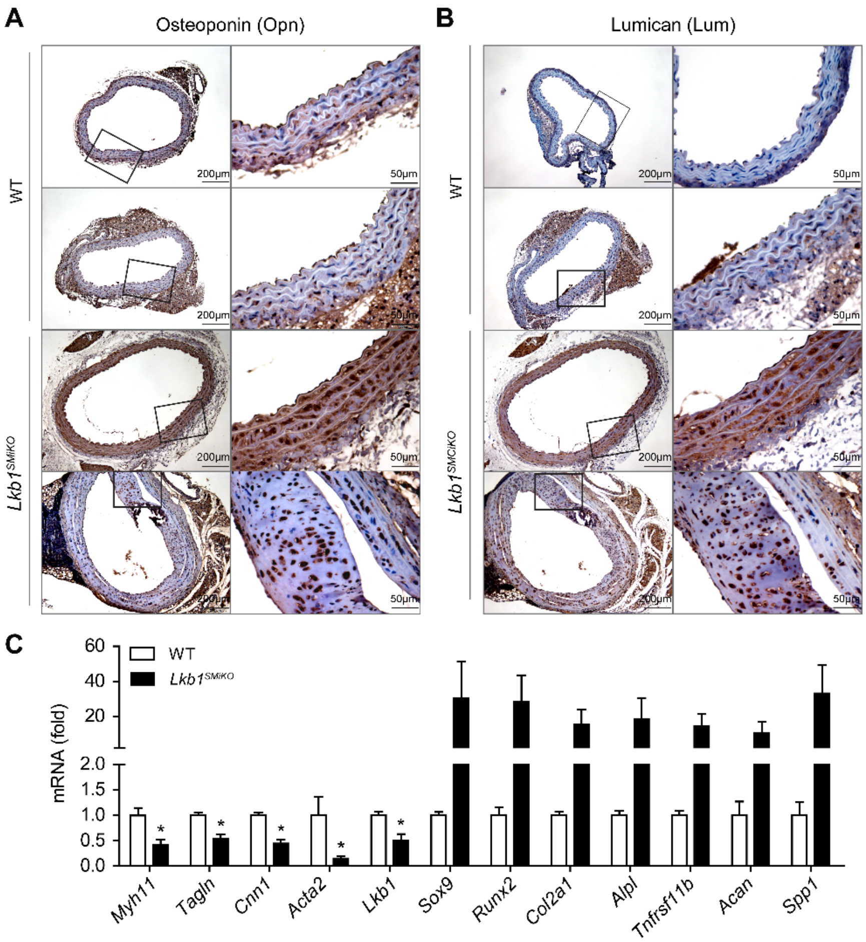


**Figure S11. Chondrocyte- and fibroblast-associated genes were increased in the aortas of *Lkb1*$^{SMiKO}$ mice**

(A and B) Immunohistochemical analysis of osteopontin (Opn) and lumican (Lum) in the thoracic aortas of WT and *Lkb1*$^{SMiKO}$ mice at 6 months post-TAM induction.

(C) Real-time PCR analysis of the indicated genes in the aortas of WT and *Lkb1*$^{SMiKO}$ mice at 6 months post-TAM induction.

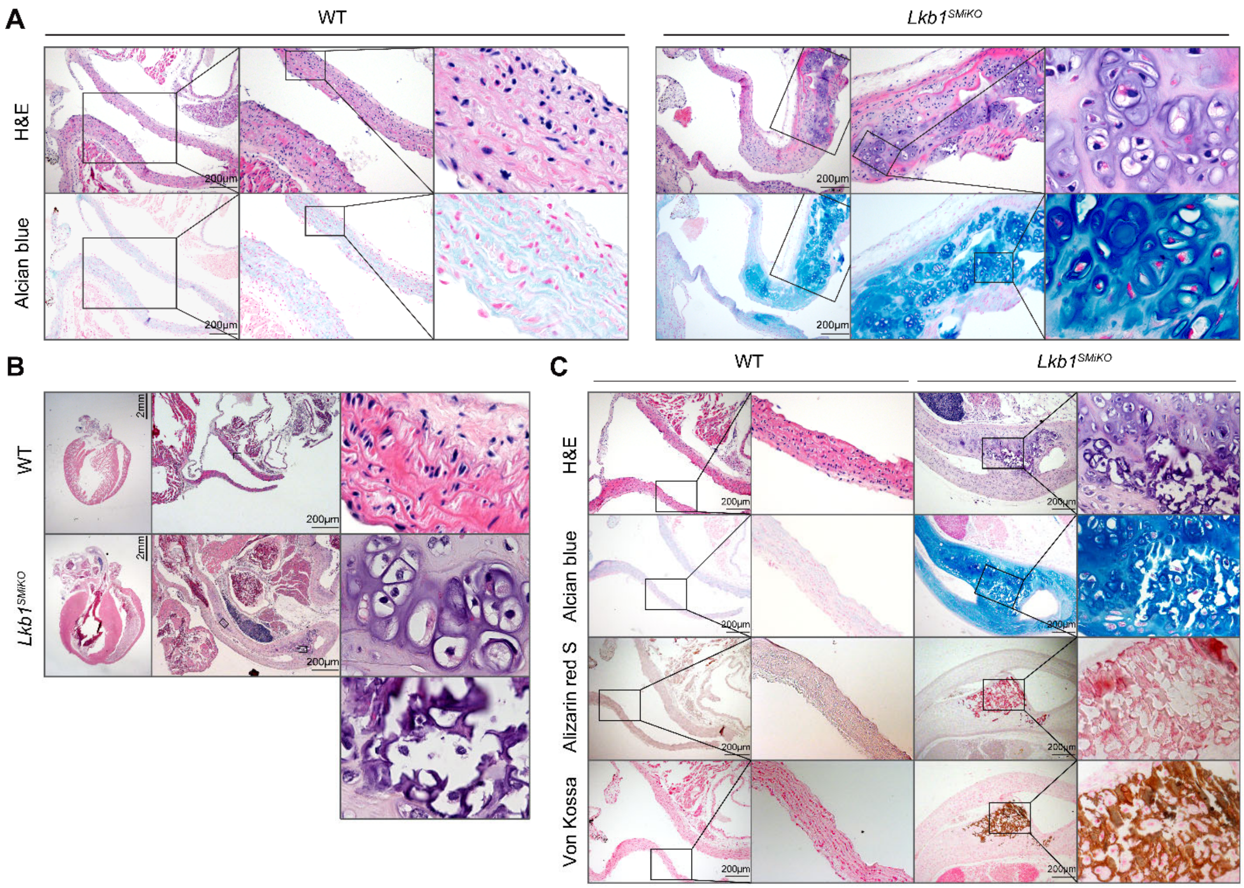


**Figure S12. Histological analysis of the aortic roots from WT and *Lkb1$^{SMiKO}$* mice.**

(A) H&E and Alcian blue staining of the aortic roots from WT and *Lkb1$^{SMiKO}$* mice at 4.5 months post-TAM induction.

(B and C) H&E, Alcian blue, Alizarin red S, and Von Kossa staining of the aortic roots from WT and *Lkb1$^{SMiKO}$* mice at 6 months post-TAM induction.

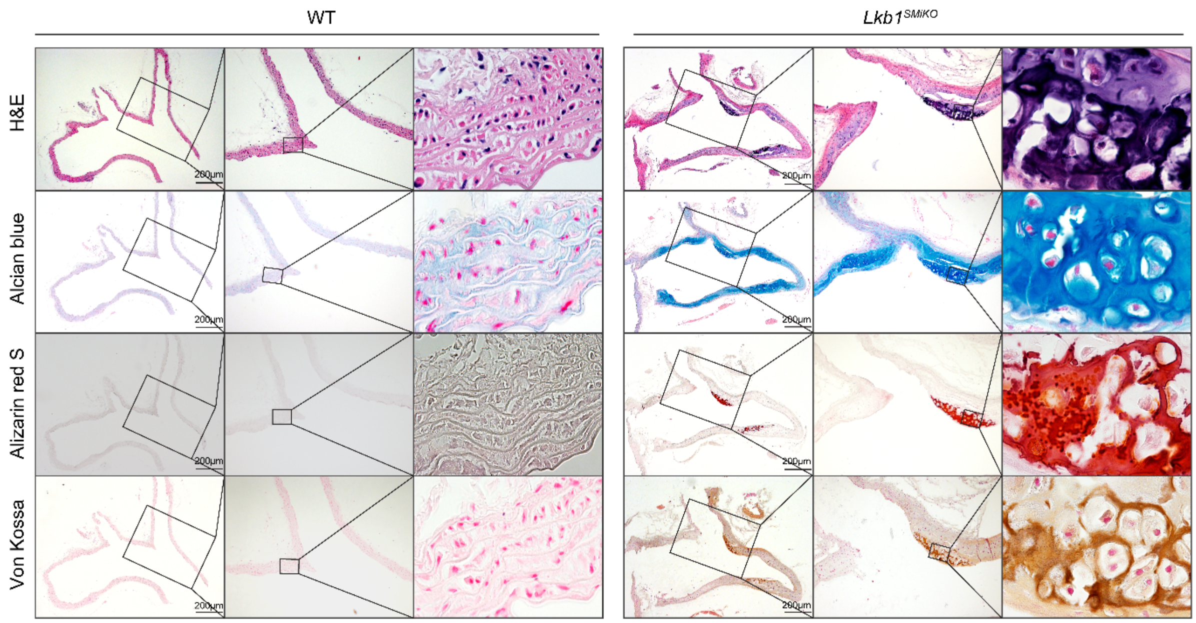


**Figure S13. Histological analysis of the aortic arches from WT and *Lkb1*$^{SMiKO}$ mice**

H&E, Alcian blue, Alizarin red S, and Von Kossa staining of the aortic arches from WT and *Lkb1*$^{SMiKO}$ mice at 4.5 months post-TAM induction.

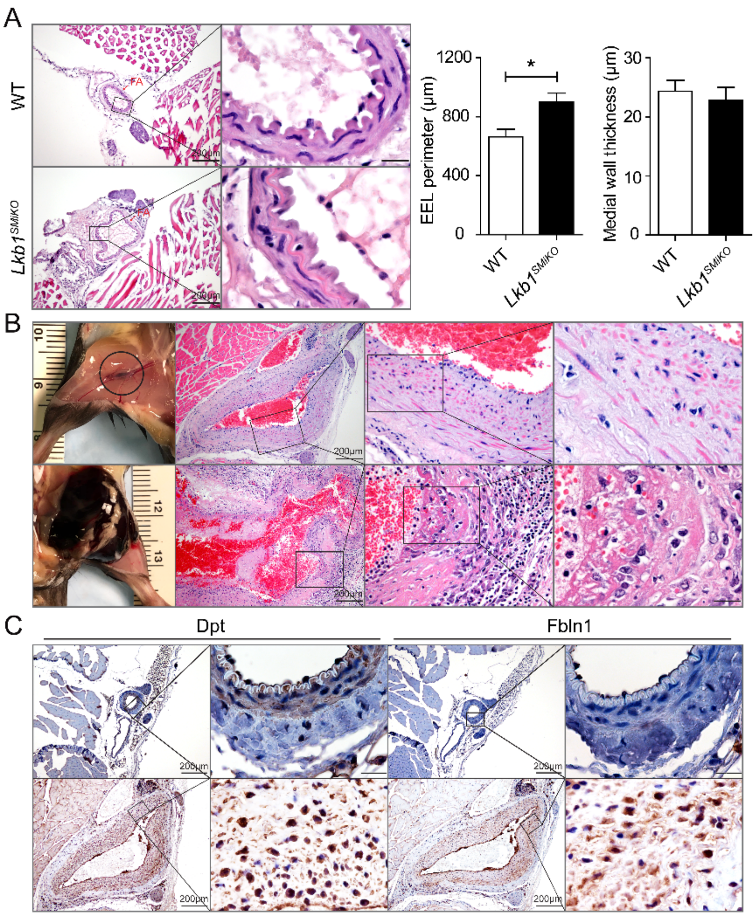


**Figure S14. Histological analysis of the femoral arteries of WT and *Lkb1*$^{SMiKO}$ mice**

(A) Representative images of H&E staining and quantification of the external elastic lamina (EEL) perimeter and medial wall thickness of the femoral arteries of WT and *Lkb1*$^{SMiKO}$ mice at 1.0-1.5 months post-TAM induction.

(B) Macroscopic images and H&E staining of non-ruptured and ruptured femoral artery aneurysms from *Lkb1*$^{SMCiKO}$ mice.

(C) Immunohistochemical analysis of dermatopontin (Dpt) and fibulin 1 (Fbln1) in the femoral arteries of *Lkb1*$^{SMiKO}$ mice.

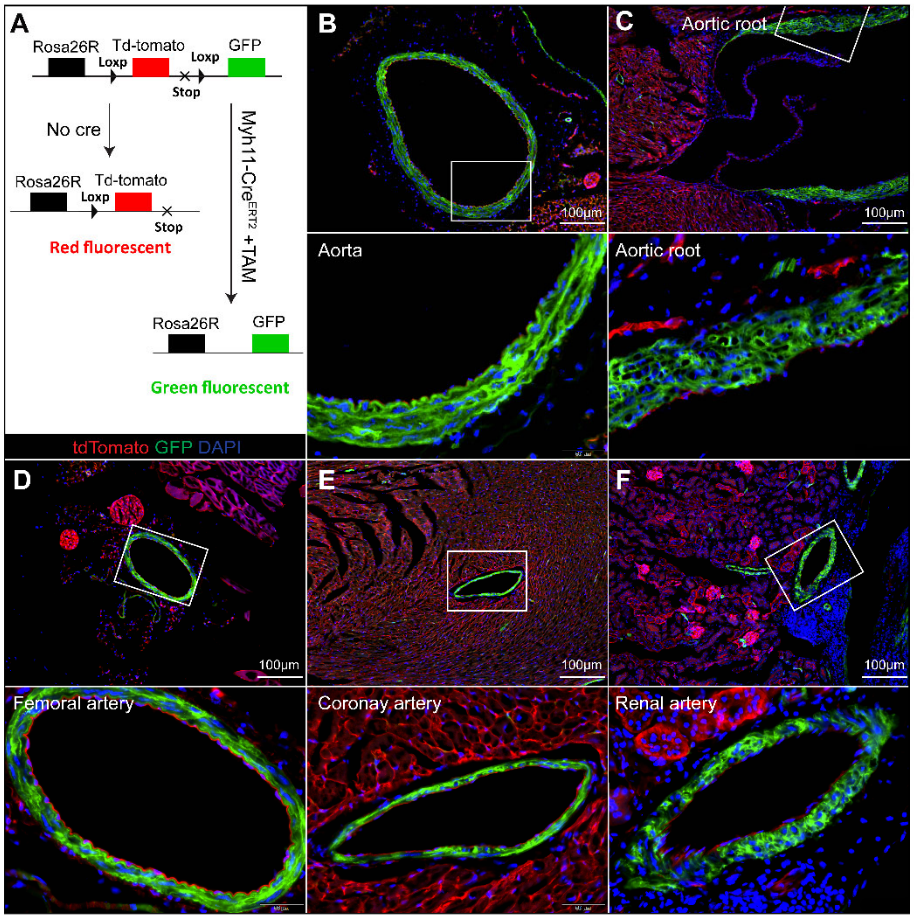


**Figure S15. Generation of lineage-tracing in *Lkb1$^{flox/flox}$*;*Myh11-Cre/ERT2*;*ROSA$^{mT/mG}$* mice.** (A) Schematic of experiments on lineage-tracing *Lkb1$^{flox/flox}$*;*Myh11-Cre/ERT2*;*ROSA$^{mT/mG}$* mice. (B to F) Immunofluorescence images of different tissue sections of *Lkb1$^{flox/flox}$*;*Myh11-Cre/ERT2*;*ROSA$^{mT/mG}$* mice, including abdominal aorta (B), aortic root (C), femoral artery (D), coronary artery (E), and renal artery (F).

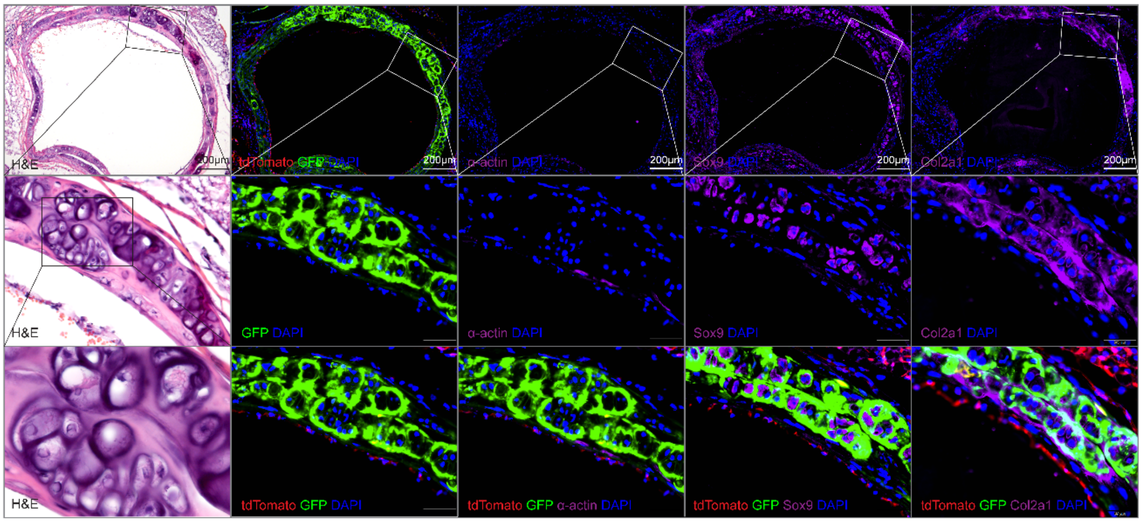


**Figure S16. Lineage tracing of VSMC in *Lkb1*$^{SMiKO}$ mice.**

H&E staining and immunostaining of α-actin, Sox9, or Col2a1 in abdominal aortas from *Lkb1*$^{+/+}$;*Myh11-Cre/ERT2*;*ROSA*$^{mT/}$*mG* and *Lkb1*$^{fl/fl}$;*Myh11-Cre/ERT2*;*ROSA*$^{mT/mG}$ mice at 6.0 months post-TAM induction.

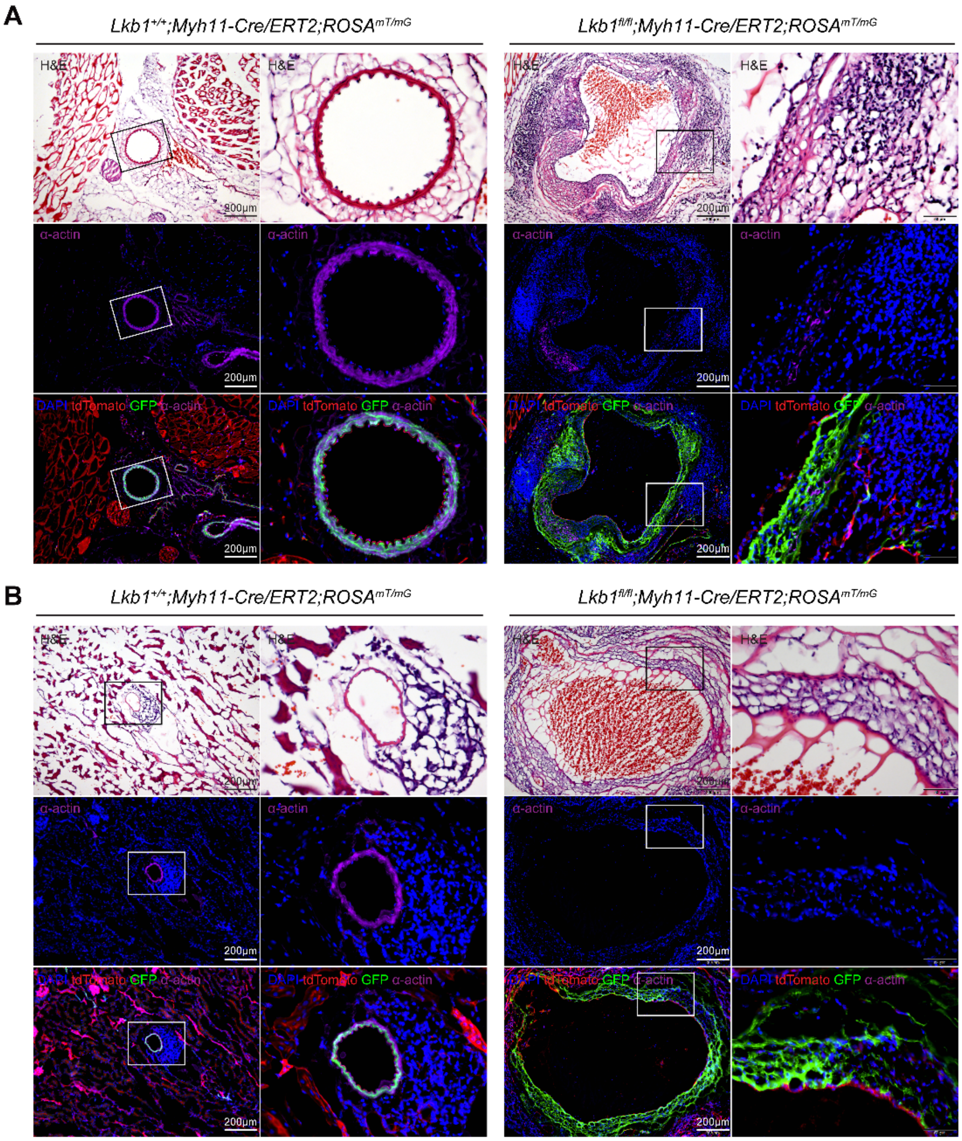


**Figure S17. Lineage tracing of VSMC in *Lkb1*$^{SMiKO}$ mice.**

**(A-B)** H&E staining and ant–α-actin co-immunostaining (purple) of femoral (A) and renal (B) arteries of *Lkb1*$^{+/+}$;*Myh11-Cre/ERT2*;*ROSA*$^{mT/}$*mG* and *Lkb1*$^{fl/fl}$;*Myh11-Cre/ERT2*;*ROSA*$^{mT/mG}$ mice at 6.0 months post-TAM induction.

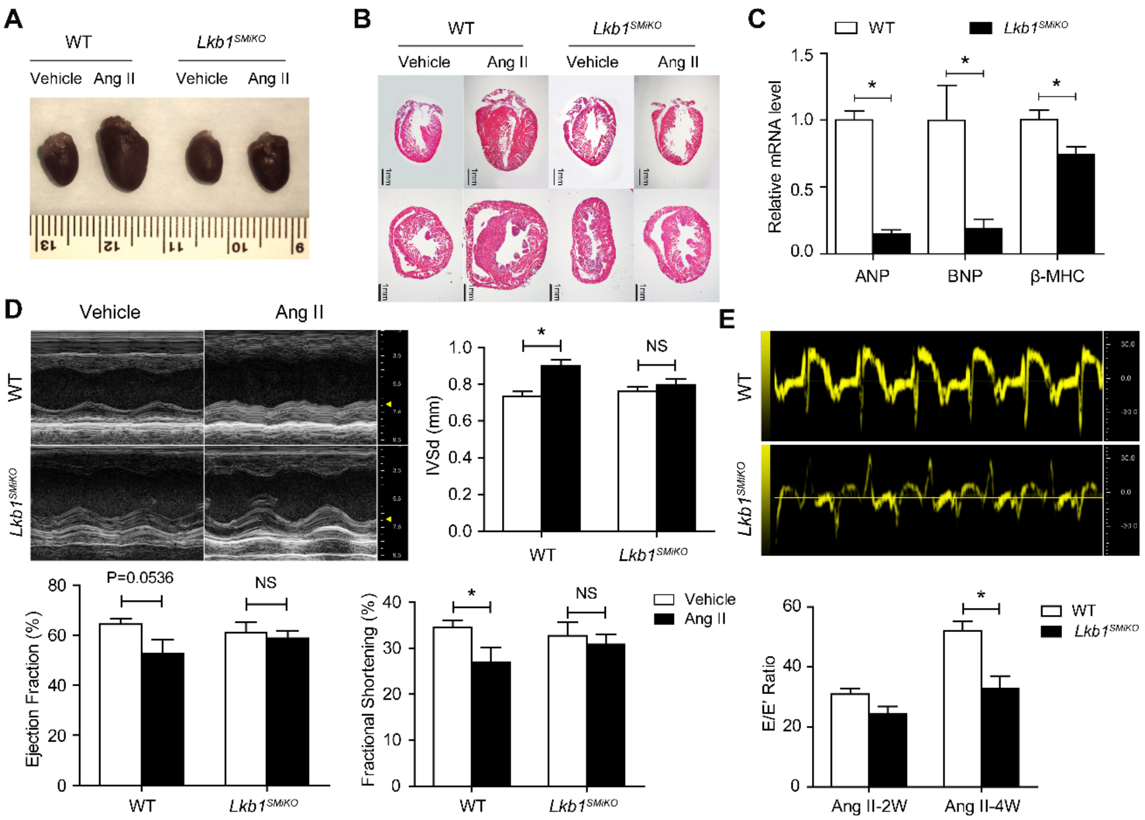


**Figure S18. SMC-specific ablation of Lkb1 abolished Ang II-induced cardiac hypertrophy**

(A and B) Representative macroscopic and H&E-stained images of the heart sections of WT and *Lkb1$^{SMiKO}$* mice infused with vehicle or AngII.

(C) ANP, BNP, and β-MHC mRNA expression.

(D and E) Heart echocardiography measurements (ejection fraction, fractional shortening, IVS, and E/E' ratio) from WT and *Lkb1$^{SMiKO}$* mice treated with vehicle or AngII. IVS, interventricular septum.

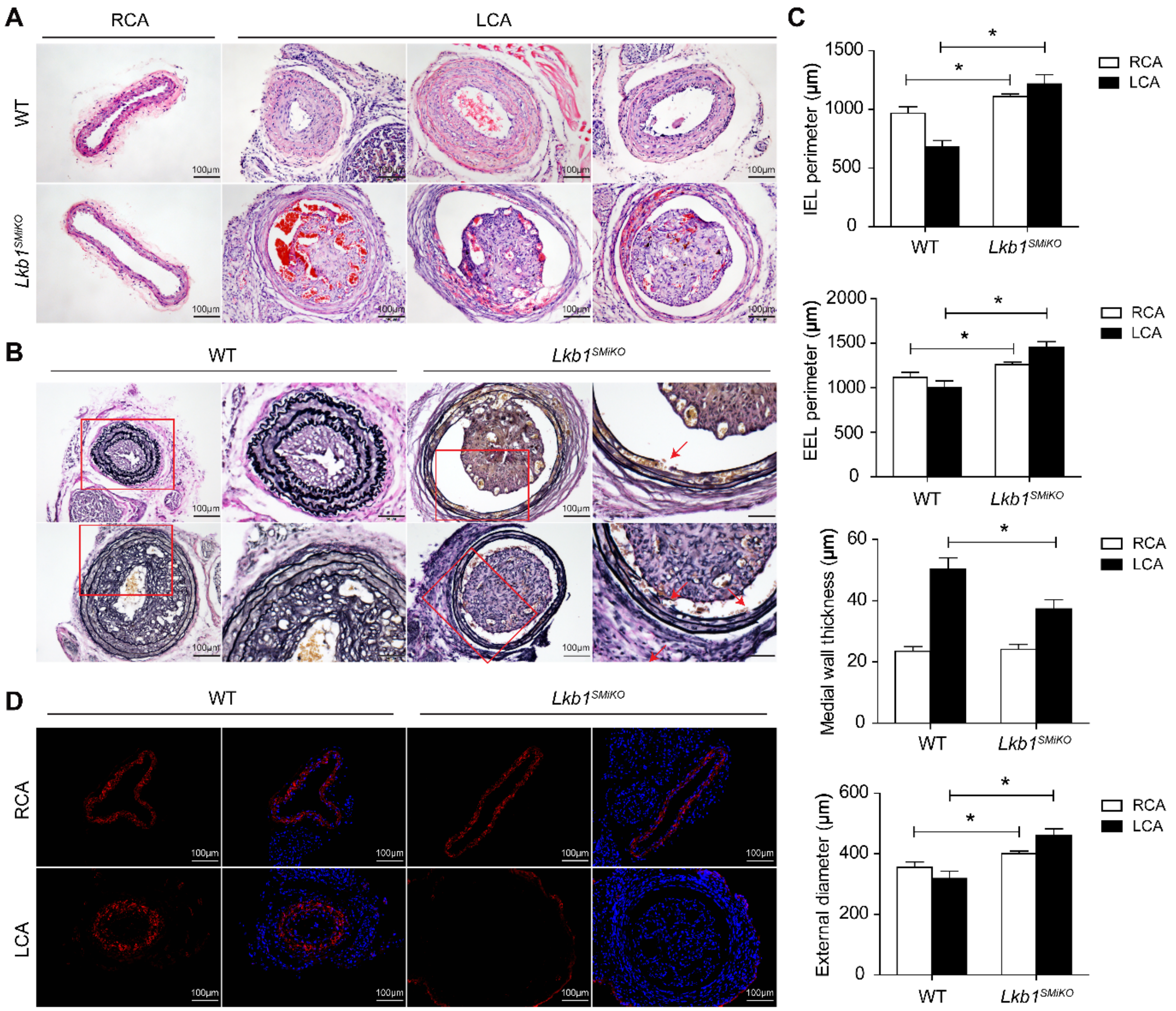


**Figure S19. SMC-specific ablation of *Lkb1* abolished injury-induced neointima formation**

**(A)** Representative images of H&E staining of the ligated (LCA) or contralateral (RCA) carotid arteries of WT and *Lkb1*$^{SMCiKO}$ mice.

**(B)** Representative images of elastin staining of the ligated or contralateral carotid arteries of WT and *Lkb1*$^{SMCiKO}$ mice. Red arrows denote disruption of the elastic lamina.

**(C)** Quantification of the internal and external elastic laminae (IEL and EEL, respectively) perimeters, medial wall thickness, and outer diameter of ligated or contralateral carotid arteries of WT and *Lkb1*$^{SMCiKO}$ mice.

**(D)** IF staining of α-actin of ligated or contralateral carotid arteries of WT and *Lkb1*$^{SMCiKO}$ mice.

**Table S1. The number of homozygous embryos at different stages of development**

| Time point | Mating mice | No. of embryos | No. of homozygous embryos | | |
|---|---|---|---|---|---|
| | | | Expected | Observed | Visibly abnormal |
| E8.5 | $Lkb1^{fl/+};Cre^{tg/0}$ × $Lkb1^{fl/+};Cre^{tg/0}$ | 15 | 2.8125 | 4 | 0 |
| E9.5 | $Lkb1^{fl/+};Cre^{tg/tg}$ × $Lkb1^{fl/+};Cre^{tg/tg}$ | 8 | 2 | 5 | 4 |
| E11.5 | $Lkb1^{fl/fl};Cre^{0/0}$ × $Lkb1^{fl/+};Cre^{tg/tg}$ | 11 | 5.5 | 3 | 3 |
| E12.5 | $Lkb1^{fl/+};Cre^{tg/tg}$ × $Lkb1^{fl/fl};Cre^{0/0}$ | 10 | 5 | 2 | 2 |
| E14.5 | $Lkb1^{fl/+};Cre^{tg/0}$ × $Lkb1^{fl/+};Cre^{tg/0}$ | 11 | 2.0625 | 4 | 4 |
| E15.5 | $Lkb1^{fl/+};Cre^{tg/0}$ × $Lkb1^{fl/+};Cre^{tg/0}$ | 9 | 1.6875 | 1 | 1 |
| E18.5 | $Lkb1^{fl/+};Cre^{tg/0}$ × $Lkb1^{fl/+};Cre^{tg/0}$ | 7 | 1.3125 | 3 | 3 |
| E20.5 | $Lkb1^{fl/+};Cre^{tg/tg}$ × $Lkb1^{fl/fl};Cre^{0/0}$ | 9 | 4.5 | 4 | 4 |
| Total | | 80 | 24.875 | 26 | 21 |

(Homozygous: $Lkb1^{flox/flox}$;$Tagln$-$Cre^{tg}$)

**Table S2. The primer sequences for qPCR in this study**

| Genes | | Sequence (5'-3') |
|---|---|---|
| *18S* | Forward | GTCTGTGATGCCCTTAGATG |
| *18S* | Reverse | AGCTTATGACCCGCACTTAC |
| *Lkb1* | Forward | GACTTCACAGTGCCTGGACA |
| *Lkb1* | Reverse | CACAAACAGCCTTGGGCAAA |
| *Ifi27l2a* | Forward | CTGTTTGGCTCTGCCATAGGAG |
| *Ifi27l2a* | Reverse | CCTAGGATGGCATTTGTTGATGTGG |
| *Rasl10b* | Forward | GACAGCTTTGAGTACGTCAAGA |
| *Rasl10b* | Reverse | TAGCCGCACTTCCAGGTCT |
| *Hapln1* | Forward | TCACACAAAGGACCAGAATCG |
| *Hapln1* | Reverse | TGGTAATCTTGAAGTCTCGAAAGG |
| *Tcap* | Forward | CGTGGGCTACAGGAATACCAG |
| *Tcap* | Reverse | GAGACATGGATCGAGACAGGG |
| *Por* | Forward | AGCACAACGGACATTGTTCTG |
| *Por* | Reverse | GCTGAACTCCGGTATCTCTTCT |
| *Acta2* | Forward | CTGACAGAGGCACCACTGAA |
| *Acta2* | Reverse | GAAATAGCCAAGCTCAG |
| *Cnn1* | Forward | GCACATTTTAACCGAGGTCC |
| *Cnn1* | Reverse | TGACCTTCTTCACAGAACCC |
| *Myh11* | Forward | TGGACACCATGTCAGGGAAA |
| *Myh11* | Reverse | ATGGACACAAGTGCTAAGCAGTCT |
| *Tagln* | Forward | CCCAGACACCGAAGCTACTC |
| *Tagln* | Reverse | TCGATCCCTCAGGATACAGG |
| *Sox9* | Forward | GAGGCCACGGAACAGACTCA |
| *Sox9* | Reverse | CAGCGCCTTGAAGATAGCATT |
| *Runx2* | Forward | CACCGAGACCAACCGAGTCA |
| *Runx2* | Reverse | TGCTCGGATCCCAAAAGAAG |
| *Col2a1* | Forward | TTAGAAAGGGGAGCACAGTCC |
| *Col2a1* | Reverse | TACACTGCCATGAAGCATGG |
| ALP | Forward | GTGCAGTCTGTGTCTTGCCTG |
| ALP | Reverse | CCTTGCCTGTATCTGGAATCCT |
| *Tnfrsf11b* | Forward | AGAGCAAACCTTCCAGCTGC |
| *Tnfrsf11b* | Reverse | CTGCTCTGTGGTGAGGTTCG |
| *Acan* | Forward | TTCCATCTGGAGGAGAGGG |
| *Acan* | Reverse | ATCTACTCCTGAAGCAGATGTC |
| *Spp1* | Forward | CTTCCAAGCAATTCCAATGAAAG |
| *Spp1* | Reverse | TGTGTACTAGCAGTGACGG |
| *Cxcl12* | Forward | TGCATCAGTGACGGTAAACCA |
| *Cxcl12* | Reverse | TTCTTCAGCCGTGCAACAATC |
| *H2-Ab1* | Forward | ACAGCTTATTAGGAATGGGGACT |
| *H2-Ab1* | Reverse | CACGGTGATGGGACTCTTCA |
| *C3* | Forward | CCAGCTCCCCATTAGCTCTG |
| *C3* | Reverse | GCACTTGCCTCTTTAGGAAGTC |

| | | |
|---|---|---|
| *Cd74* | Forward | CCGCCTAGACAAGCTGACC |
| *Cd74* | Reverse | ACAGGTTTGGCAGATTTCGGA |
| *Vcam1* | Forward | AGTTGGGGATTCGGTTGTTCT |
| *Vcam1* | Reverse | CCCCTCATTCCTTACCACCC |
